# Personalized Multiscale Modeling of Left Atrial Mechanics and Blood Flow

**DOI:** 10.1101/2025.04.26.650771

**Authors:** Lei Shi, Ian Y. Chen, Vijay Vedula

## Abstract

We present a personalized multiscale mechanics model of the left atrium (LA) to simulate its deformation throughout the cardiac cycle and drive blood flow. Our patient data-driven model tightly integrates 3D structural mechanics of the LA myocardium, incorporating both passive and active components, with a 0D closed-loop lumped parameter network (LPN)-based circulatory system model. A finite element (FE) model of LA tissue is constructed from the patient’s images, assuming uniform thickness and employing rule-based fiber directions, a structurally based constitutive model for the passive mechanics, and a phenomenological contraction model while applying physiologically relevant boundary conditions. We then adopted a multi-step personalization approach, in which the LPN parameters with a surrogate LA model are first optimized to match cuff-based blood pressures and cardiac lumen volumes derived from time-resolved 3D gated computed tomography angiography (CTA) images. The surrogate LA pressure during passive expansion is used to estimate myocardial passive mechanics parameters and the reference unloaded configuration using an inverse finite element analysis (iFEA) framework. Finally, a robust multiscale coupling is applied between the iFEA-optimized FE model and the tuned 0D LPN model to characterize LA contraction. This effectively captures the 8-shaped pressure-volume curve and reasonably aligns with the image-based cavity volumes and deformation. The resulting simulation-predicted deformation is imposed as a moving-wall boundary condition to model atrial hemodynamics. Overall, this comprehensive digital twinning platform could be applied to study LA biomechanics in health and disease and assist in devising personalized treatment plans.

## 1. Introduction

Computational modeling is increasingly becoming an indispensable tool for studying cardiovascular disease and function [1, 2, 3, 4]. Significant advances were made in modeling cardiac function, resolving electrical activation [5, 6, 7, 8, 9, 10], tissue contraction and expansion [11, 12, 13, 14, 15, 16, 17], and blood flow [18, 19, 20, 21, 22, 23]. Multiphysics, multi-chamber models based on geometries constructed from patient images, and realistic boundary conditions are being actively developed [24, 25, 26, 27, 28, 29, 30, 31, 32, 33, 34, 35, 36, 37, 38]. Most of these developments, however, are centered on modeling the lower pumping chambers, typically involving the left ventricle (LV) in a univentricular or a biventricular setup. Meanwhile, modeling the upper chamber, particularly the mechanics of the left atrium (LA), received less attention and is the primary focus of this work. In what follows, we will briefly review LA’s complex topology and function and the most common diseases affecting it. We will then highlight a few relevant computational studies focused on LA mechanics and blood flow before motivating the reader on the unmet need to develop a personalized multiscale model of LA mechanics.

The LA has a complex structure and function (Fig. 1) [39, 40, 41, 42, 43, 44]. The pulmonary veins are aligned asymmetrically with respect to the LA cavity. The left pulmonary veins (LPVs), connected to the left lung, are positioned lower in the atrial cavity near the mitral orifice, while the right pulmonary veins (RPVs), receiving blood from the right lung, are placed higher with respect to the LA cavity (Fig. 1A). LA is attached to a highly distensible and trabeculated sac called the left atrial appendage (LAA), positioned near the mitral orifice and sandwiched between LA and the pericardium [42]. This unique LA structure results in a chaotic flow in the LA cavity due to interacting vortices from each pulmonary vein. Further, the flow from the LPVs loops around the LA cavity toward the mitral valve (MV) annulus, whereas the RPVs eject a streamlined flow directed at the MV plane. These complex LA flow features are well characterized using time-resolved phase-contrast magnetic resonance imaging (pcMRI) and image-based blood flow simulations [41, 45].

**Figure 1:**
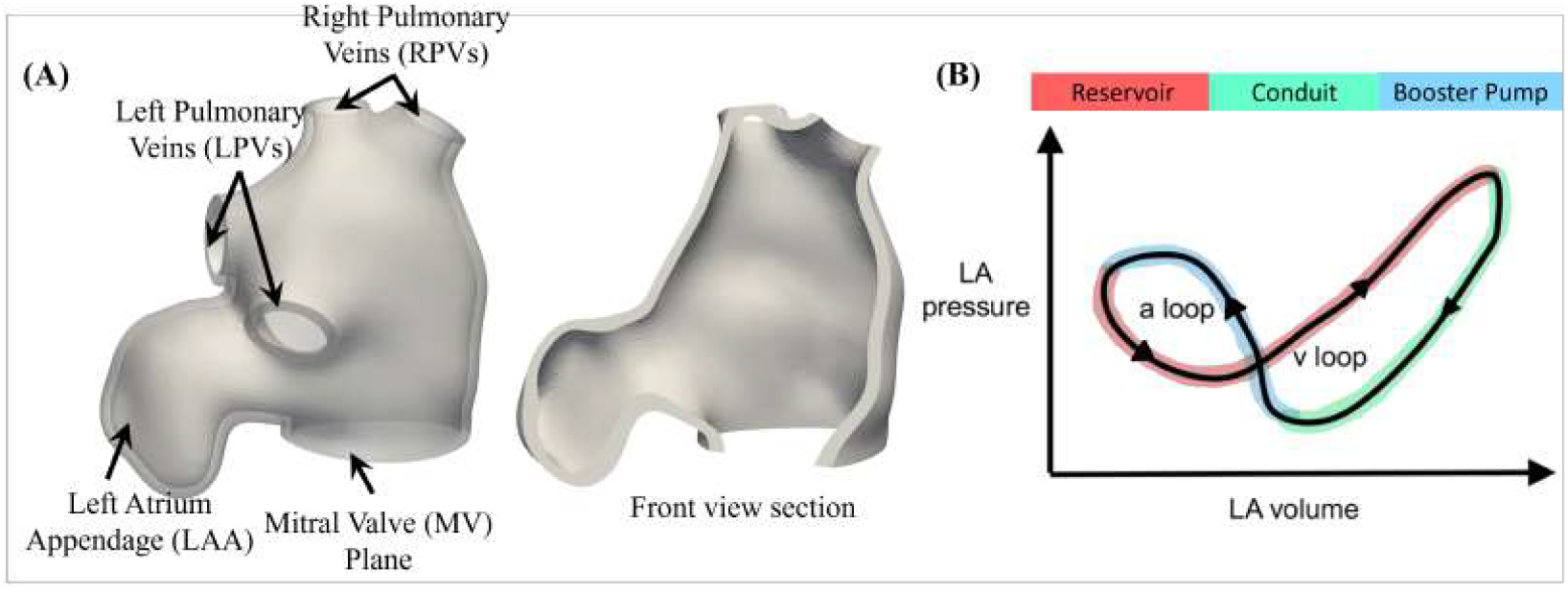
(A) The structure of the left atrium (LA). (B) A schematic of LA pressure-volume (P–V) diagram, comprising an ‘*a*–loop’ and a ‘*v* –loop’, characterized by reservoir, conduit, and booster pump function.

The LA exhibits heterogeneity in its structure, often ignored when modeling myocyte action potentials or myocardial mechanics. Regional variation in LA thickness is well known [46, 47, 48, 49, 50], although there is conflicting data on the absolute thickness values [39, 40, 46, 51]. While prior computational studies have reported that variable thickness affects local tissue mechanics [49, 50], assuming a uniform wall thickness for modeling LA electrophysiology and mechanics is common [52, 53, 54, 55, 56]. Likewise, although four pulmonary vein attachments (two LPVs and two RPVs) with the LA are routinely observed in patients, several other normal variations exist [57, 58, 59], the biomechanical effects of which have not received much attention until recently [59, 60].

The complexity of the LA structure is complemented by its three-fold function during the cardiac cycle (Fig. 1B), comprising a reservoir phase, a conduit phase, and a booster pump phase [42, 43, 52, 61]. During ventricular ejection (systole), the mitral valve is closed, and the LA forms a reservoir of blood flowing in from the pulmonary veins. During early ventricular filling (E-wave), the low pressure in the ventricular cavity drives blood from the LA, acting as a conduit passage. The atrial filling (A-wave) follows when the LA acts as a booster pump, forcefully delivering blood into the LV. This complex three-fold LA function, with concomitant changes in LA pressure and volume, translates into a lazy eight-shaped (lemniscate) loop on the pressure-volume (P–V) chart with a distinct *a*–loop, dominated by the booster pump function, and a *v* –loop, dominated by the reservoir and conduit phases (Fig. 1B) [44, 52, 61, 62]. It is, therefore, vital to capture this physiological behavior when modeling LA mechanics.

LA is often implicated in rhythm disorders such as atrial fibrillation (AF) [63, 64], atrial flutter (AFl) [65], and atrial tachycardia (AT) [66], among which AF is the most prevalent [64], with a high potential for blood clots and stroke [67, 68, 69]. Based on 2016 estimates, over 46 million people live with AF worldwide, which is expected to grow 2.5 times in the next few decades [70]. AF is estimated to cause up to a 4-5 fold increased risk of stroke, which remains the second leading cause (∼ 17%) of all cardiovascular disease-related deaths [71]. Atrial myocardial remodeling and fibrosis play a prominent role in the occurrence and sustenance of AF [67, 72, 73]. AF manifests as an abnormal electrical activation during which the LA wall quivers and does not undergo coordinated contraction. Failing to produce the atrial kick (A-wave) impairs LA’s booster and reservoir functions, depriving the LV of an additional source of blood [62]. As a result, the ventricle undergoes compensatory remodeling, eventually leading to heart failure. This abnormal contraction pattern during AF alters LA hemodynamics, particularly in the LAA. As there is no other outlet in the LAA, blood is not effectively washed out of the LAA cavity during AF, leading to a stagnant flow that triggers clot formation [42]. When a sinus rhythm sets in following AF, the complex blood-clot interactions in the LAA could break the clot into emboli, potentially leading to stroke.

Image-based computational models were developed to study arrhythmogenesis and blood dynamics leading to clotting in patients with AF. Boyle et al. [6] used late gadolinium-enhanced magnetic resonance imaging (LGE-MRI) data to construct fibrotic LA models and simulated arrhythmia by modifying generic electrophysiology model parameters to identify optimal ablation targets and guide treatment planning [6]. On the other hand, blood flow models in LA are designed to study thrombus formation in LAA under AF conditions by imposing wall motion derived from medical image data, including computed tomography angiography (CTA) or MRI [41, 74, 75, 76, 77, 78, 79, 80]. In this approach, LA models are constructed by segmenting the time-dependent image data acquired during AF or sinus rhythm, followed by image registration or point cloud deformation techniques to extract the LA wall motion. By imposing this wall motion as a moving-wall boundary condition, changes to blood flow and related hemodynamic measures are analyzed for flow stasis and thrombogenicity [74, 75, 76, 77, 81]. Thrombosis is modeled using a simplified or comprehensive coagulation cascade model to evaluate the likelihood of clot formation in the low-velocity regions, such as the LAA cavity. Flow boundary conditions are applied by imposing pressures or flowrates at the pulmonary veins and MV plane, often derived from echo or 4D-flow MRI data.

Although image-based computational models have been insightful in quantifying abnormal blood flow patterns and in identifying potential flow-mediated thrombogenic biomarkers [55, 75, 81], limitations remain as they lack predictive capability by relying on *status quo* acquired image data. **A personalized multiscale multiphysics model of LA provides a comprehensive end-to-end platform to study AF-related stroke and develop personalized treatment plans**. This includes coupling atrial myocyte excitation and electrical conduction with parameters matching electrocardiogram (ECG) readings, personalized tissue mechanics matching image-based deformation, and multiscale blood flow models tuned to align with the patient’s clinically measured hemodynamic data. However, studies involving electromechanical models of LA were mostly limited to understanding the effects of thickness and fiber directions on wall mechanics [49, 50], or were developed as part of four-chamber or left-heart models that involve several underlying simplifications, such as employing generic material parameters and simplified boundary conditions [9, 35, 37, 52, 82, 83, 84]. These simplifications limit our ability to perform patient-specific modeling and use simulations to develop targeted treatment plans. Nevertheless, novel image-derived multiphysics modeling approaches have been developed to bridge the gap between traditional atrial electromechanics and blood flow modeling communities [56], and our work aims to make a seminal contribution in this realm.

In this study, we have developed a multiscale computational modeling framework to simulate atrial myocardial mechanics, with the model parameters personalized to match the patient data. The resulting endocardial motion is used to drive blood flow through the LA cavity under personalized hemodynamic boundary conditions. Our framework relies on a patient’s time-resolved 3D gated-CTA images and clinical measurements, including cuff-based pressures and data from ECG waveforms. A combination of manual segmentation, point cloud registration, and neural network-based automatic segmentation and mesh tracking methods is used to extract the LA blood pool, track its motion, and measure the changes in the cavity volumes over the cardiac cycle [85]. We then adopted a multi-step personalization strategy in which we first tuned a fully 0D lumped parameter network (LPN) model, incorporating a surrogate model of LA, to match clinical measurements. We then applied our recently developed inverse finite element analysis (iFEA) framework to characterize the LA myocardial passive mechanics, using the pressure data from the optimized 0D model, while simultaneously estimating the unloaded reference configuration [85]. Subsequently, multiscale mechanics simulations are performed by iterating the contraction model parameters to closely align the predicted cavity volumes and deformation patterns with the image data over the cardiac cycle. These multiscale simulations employ a modular implicit coupling between the 3D finite element (FE) model in its reference unloaded state and the optimized 0D LPN network [86]. Finally, blood flow through the LA cavity is simulated by imposing the LA endocardial wall motion as a moving-wall boundary condition, augmented with pulmonary pressures and mitral valve flow rate. The wall motion and flow boundary conditions are retrieved from the personalized multiscale mechanics simulations.

The manuscript is structured as follows: Section 2 outlines our workflow, starting with describing the multiscale mechanics model of LA, including methods adopted to construct the 3D FE model of LA myocardium from patient data, a brief overview of myocardial mechanics formulation, and the multiscale coupling between the 3D FE model and the 0D LPN model of blood circulation. We then described the multi-step personalization process, which includes estimating LPN parameters, the iFEA pipeline to characterize passive mechanics, and estimating the contraction model parameters. We then provided a quick overview of our moving-domain blood flow formulation to simulate atrial hemodynamics before providing the results in Section 3. In Section 4, we discuss the results, provide a rationale for the various modeling decisions made while creating the entire pipeline, followed by an analysis of the effects of mesh size, peak active stress, and fiber direction parameters on the LA mechanics. Finally, we acknowledge limitations of the current work, provide future directions, and conclude.

## 2. Methods

We developed a comprehensive workflow for patient-specific modeling of LA mechanics (Fig. 2). Our workflow has three major components: (i) extracting all the required and available patient data (Fig. 2, left), (ii) creating a multiscale model of LA mechanics (Fig. 2, center), and (iii) estimating parameters of the multiscale FEA model to simulate LA myocardial deformation throughout the cardiac cycle that aligns reasonably with the clinical data (Fig. 2, right). We will provide a detailed description of these components in the following subsections.

**Figure 2:**
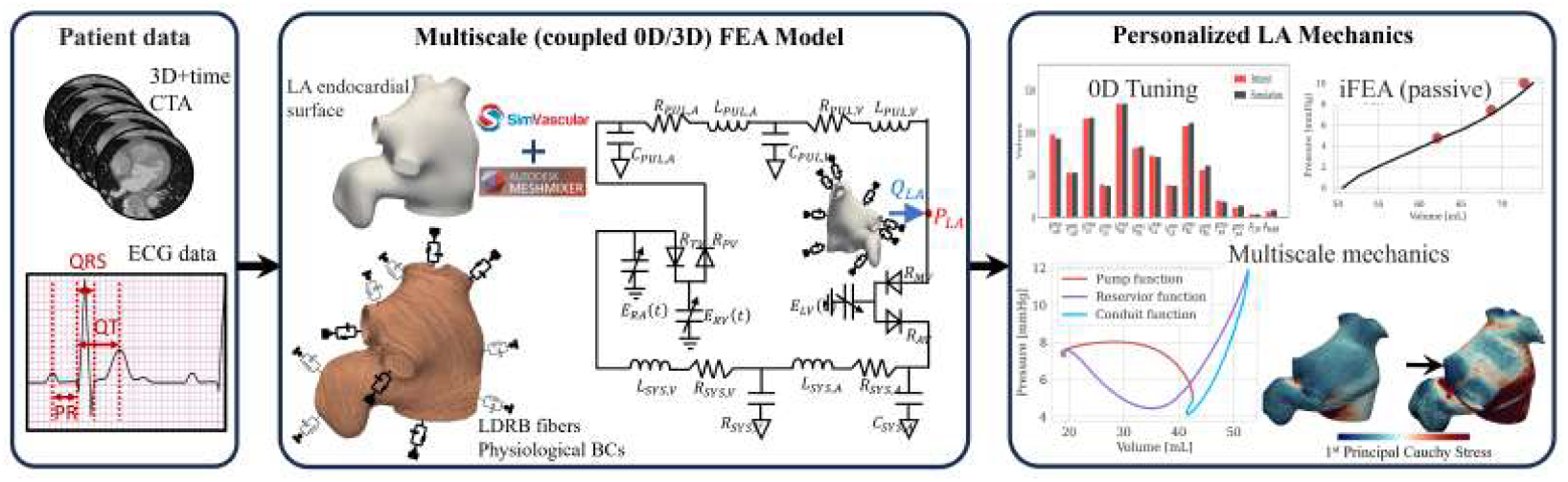
Workflow to simulate patient-specific LA mechanics. (left) The workflow begins with acquiring patient data, including time-dependent gated computed tomography angiography (CTA) images, with accompanying electrocardiogram (ECG) and cuff-based pressure measurements. (center) A multiscale finite element analysis (FEA) model is created by coupling the 3D finite element (FE) model with a 0D lumped parameter network (LPN) model of the circulatory system [86, 87]. The 3D FE model is created using our previously developed segmentation pipeline [85], incorporating detailed myocardial structure and rule-based fiber directions [85, 88], and applying physiological boundary conditions [32, 85]. (right) A multi-step tuning procedure is adopted to estimate the multiscale FEA model parameters to personalize myocardial mechanics throughout the cardiac cycle.

### 2.1. Patient data

A 50-year-old male subject is enrolled in this study, who was diagnosed with coronary artery disease under a stress test on echocardiography and underwent ECG-gated CTA with maximum tube current at the Veteran Affairs Palo Alto Healthcare System. Based on the echo readings, the heart chambers were determined to be normal in size without any wall motion abnormalities. Data acquisition was conducted with prior approval from the Institutional Review Board (IRB), and HIPAA-compliant procedures were followed throughout the study. More details about the imaging modality and CTA protocol are provided in Chen et al. [89]. These CTA images have a high spatial resolution (512 ×512 213 ×voxels) with an individual voxel size of 0.39× 0.39 ×0.7mm^3^. About 20 volumes are captured through the cardiac cycle, measured between successive R-waves on the QRS complex, uniformly sampled at every 5% R-R. Clinical measurements, including cuff-based blood pressure, ECG data, and heart rate, are also obtained (Table 1).

**Table 1:**
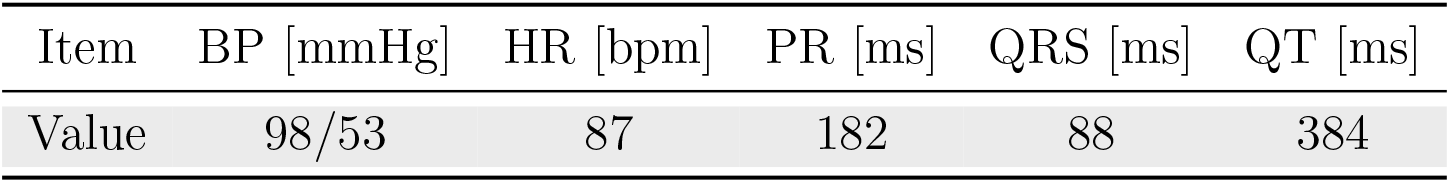
Clinical measurements of the patient used in this study. BP: blood pressure in mmHg; HR: heart rate in beats per minute (bpm); ECG: electrocardiogram; PR: the duration from P-wave to R-wave on ECG; QRS: duration of the QRS complex; QT: the duration from Q-wave to T-wave on ECG.

### 2.2. Multiscale model of LA mechanics

We developed a multiscale model of LA mechanics, coupling a 3D FE model of LA with a 0D LPN model of the circulatory system. The 3D FE model is created using our recently published segmentation pipeline [85], adopting rule-based fiber directions [85, 88], structurally-based constitutive model [85], and physiological boundary conditions [32, 85], while the 0D LPN model is adapted from Regazzoni et al. [87]. These features of the multiscale LA model, along with the solution procedure, are described briefly in the following subsections.

#### 2.2.1. 3D model segmentation

Our objective here is three-fold: (i) First, to create a 3D model of LA myocardium at its relaxed state during ventricular systole (reservoir phase, 20% R-R interval), with rule-based fiber directions, which will be used as the starting point for characterizing LA passive mechanics^*^ [85]; (ii) Second, to segment the LA endocardial surface at all phases of the cardiac cycle; and (iii) Third, to extract the endocardial cavity volumes and surface nodal displacements, to be used as target criteria during personalization procedure. We employed our previously developed manual segmentation workflow to extract the LA blood pool from 3D CTA images at 20% R-R phase [85], using SimVascular^†^ and Meshmixer^‡^. The manual approach involves creating paths and 2D segmentations and lofting the endocardial surface in SimVascular [90], following which surface smoothing and decimation in Meshmixer (Autodesk Inc.) refine the triangulated surface. The endocardial surface is then extruded along its surface normal, assuming a uniform thickness (2 mm) [40, 88, 91]. Any surface irregularities during extrusion are smoothed in Meshmixer (Autodesk Inc.), resulting in a smooth representation of LA myocardium for subsequent analysis (Fig. 1A).

Following the procedure outlined in Piersanti et al. [88], we employed a rule-based method to define fiber directions on the LA myocardium model (Fig. 3A). Specific threshold values for the LPV, the RPV, and the mitral valve annulus are *τ*_lpv_ = 0.65, *τ*_rpv_ = 0.1, and *τ*_mv_ = 0.65, respectively [85, 88]. We will assess the model’s sensitivity to these parameters in Section 4.

**Figure 3:**
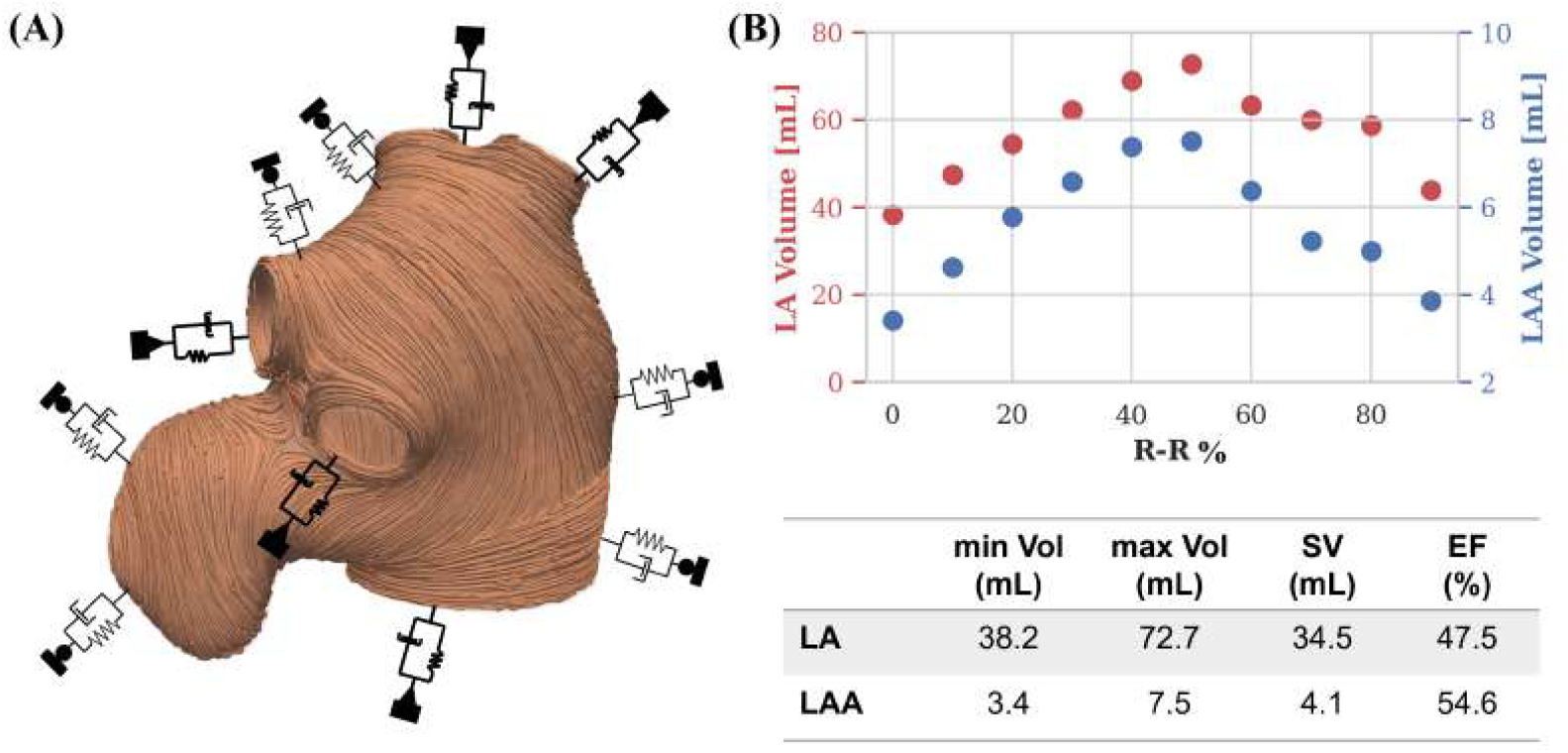
(A) The LA myocardium is segmented from CTA images during early systole (∼ 20% R-R interval) using a recently developed workflow [85]. A rule-based method is used to create LA myocardial fibers [88]. Robin boundary conditions are applied on the LA tissue while hemodynamic pressure is applied on the LA endocardium [85]. (B) (top) Changes in the cavity volumes of LA and LAA extracted from the segmented CTA images, and (bottom) relevant metrics. LA: left atrium; LAA: left atrial appendage; min/max Vol: minimum/maximum volume; SV: stroke volume; EF: ejection fraction.

We applied the above manual segmentation pipeline to extract LA endocardial surfaces at all the other phases of the cardiac cycle (0–100% R-R). We then measured the LA and LAA cavity volume changes during the cycle (Fig. 3B). Further, we also applied a Bayesian Coherent Point Drift (BCPD) algorithm [92], implemented in the Python-based open-sourced tool, *probreg*^§^, to extract endocardial nodal displacements relative to the model segmented initially (20% R-R interval). This point cloud mapping method is employed because the surface mesh topology, including nodes and element connectivity, is altered during segmentations, making it challenging to establish a nodal correspondence between successive segmentations [41, 85]. These cavity volumes (both LA and LAA) and nodal displacements at select landmarks (Fig. 5) will be used as target criteria in the objective function for estimating the multiscale model parameters (Section 2.3).

**Figure 4:**
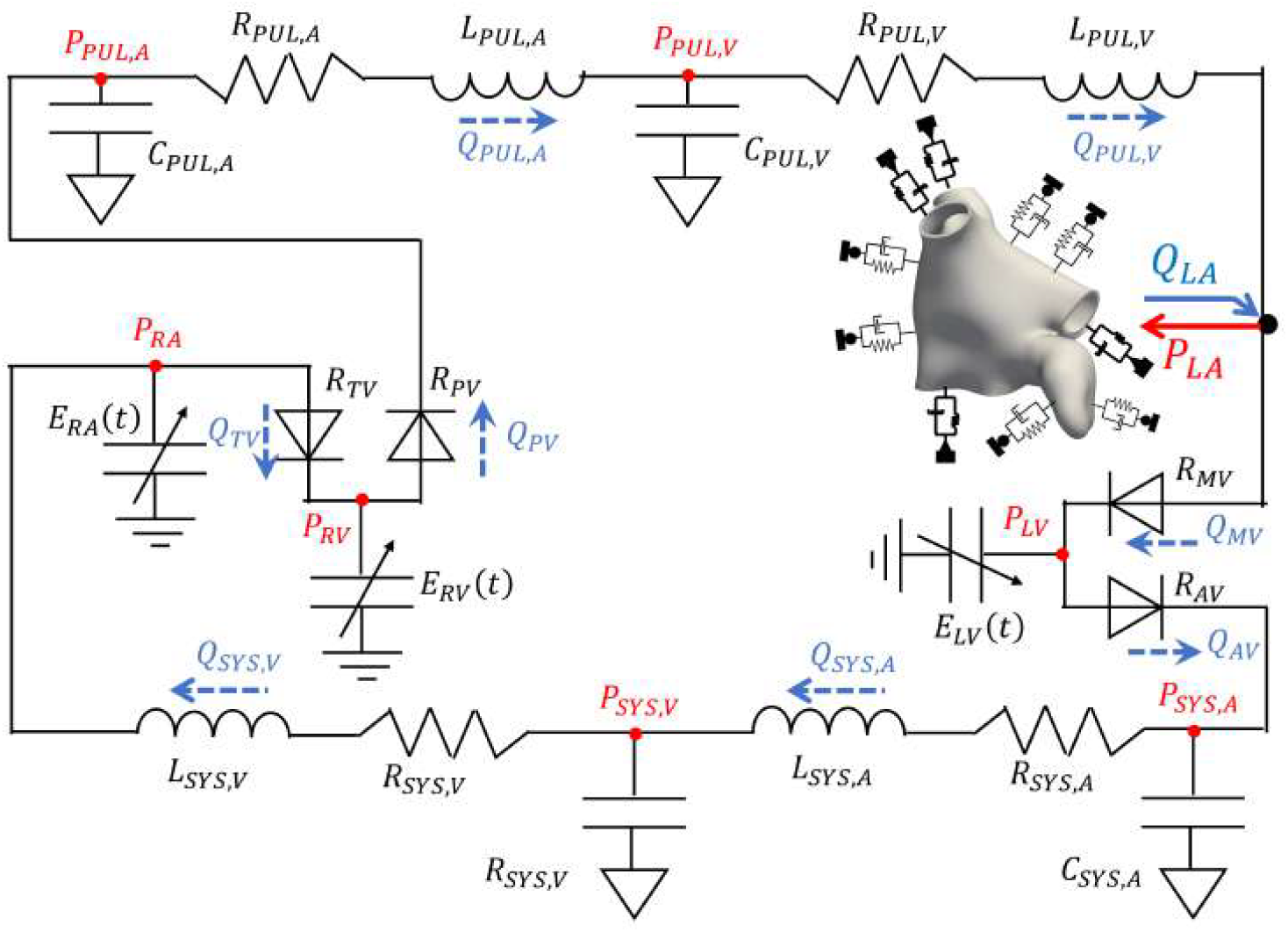
A multiscale model of LA mechanics is developed by coupling the 3D FE model of the LA myocardium with a 0D LPN-based model of the blood circulatory system, exchanging rate of volume change (*Q*_*LA*_) and pressure (*P*_*LA*_) during every nonlinear iteration until convergence [86, 87]. The parameters of the LPN model and the equations governing it are described in Appendix A. LA: left atrium; FE: finite element; LPN: lumped parameter network; *Q*_(·)_: flowrates; *P*_(·)_: pressures; *R*_(·)_: resistance; *C*_(·)_: capacitance; *L*_(·)_: inductance; *E*_(·)_: elastance; LV: left ventricle; SYS,A: systemic arteries; SYS,V: systemic veins; PUL,A: pulmonary arteries; PUL,V: pulmonary veins; RA: right atrium; RV: right ventricle; TV: tricuspid valve; PV: pulmonary valve; MV: mitral valve; AV: aortic valve.

**Figure 5:**
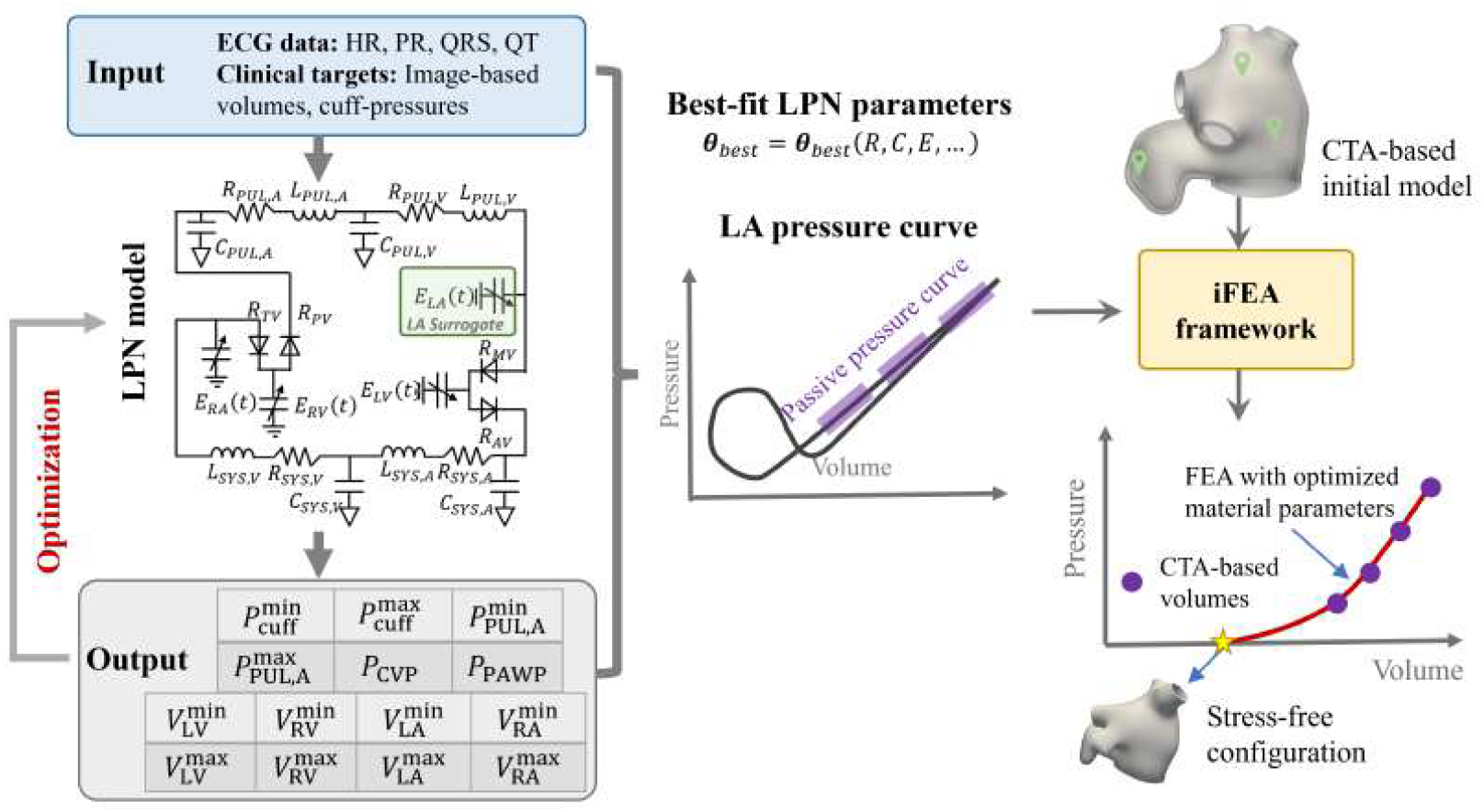
A multi-step tuning procedure is adopted to estimate the parameters of the multiscale mechanics model of LA. (left) A surrogate model of LA is used to create a fully 0D LPN model of the circulatory system. With the patient’s ECG data as input, the LPN parameters are estimated using the `L-BFGS-B` minimization algorithm to match clinical measurements, including image-based chamber volumes and available pressure data. (center-right) The LPN-predicted LA pressure during the passive expansion (reservoir phase) is extracted for estimating the myocardial passive mechanics parameters and the unloaded reference configuration using our recently developed iFEA algorithm [85]. Green dots denote the landmarks for the displacement difference calculation [85]. LA: left atrium; LPN: lumped parameter network; ECG: electrocardiography; iFEA: inverse finite element analysis; CTA: computed tomography angiography; HR: heart rate; CVP: central venous pressure; PAWP: pulmonary artery wedge pressure; PR, QRS, QT: ECG-based time intervals; Other LPN variables are described in Fig. 4.

#### 2.2.2. Myocardial mechanics formulation

We will formulate the myocardial mechanics problem by concisely introducing the nonlinear continuum mechanics framework, boundary conditions, and constitutive models. We will briefly describe the solution procedure employing stabilized finite element methods and provide additional details on the numerical methods and solver parameters.

##### Kinematics, equations of motion, and boundary conditions

Let **Ω**_***X***_ and **Ω**_***x***_ be bounded open sets in ℛ^3^ with Lipschitz boundaries and represent the reference and current configurations, respectively. In this study, **Ω**_***X***_ denotes the stress-free reference configuration of the myocardium while **Ω**_***x***_ represents the deformed myocardium at any instant during passive expansion. We define the boundary, **Γ**_***x***_ := **Γ**_endo_ **∪ Γ**_epi_ **∪ Γ**_lpv_ **∪ Γ**_rpv_ **∪ Γ**_mv_, as the union of the endocardium (**Γ**_endo_), epicardium (**Γ**_epi_), and the annular cross-sections of LPV (**Γ**_lpv_), RPV (**Γ**_rpv_), and mitral valve (**Γ**_mv_). The motion of the body is then characterized by a deformation map, *φ* : **Ω**_***X***_ **→ Ω**_***x***_ such that **x** = *φ* (**X**, *t*), where **x** is the current position of a material point at a time *t* that was originally at **X** in the reference configuration. The displacement, **u** and velocity, **v** of a material particle are defined as,

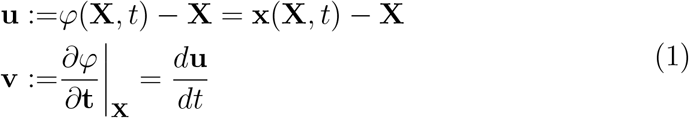

where, *d*(•)*/dt* is the total time derivative. The deformation gradient (**F**), its Jacobian determinant (*J*), the right Cauchy-Green deformation tensor (**C**), and the Green-Lagrange strain tensor (**E**) are defined as,

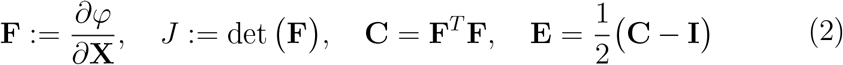

We perform a multiplicative decomposition of the deformation gradient tensor to define 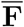 and 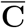 as,

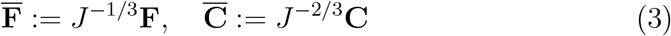

that represent the splitting of local deformation gradient (**F**) into volume-preserving (isochoric,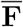) and volume-changing (dilatational, *J*^1*/*3^**I**) components. The mechanical behavior of the hyperelastic material is characterized by a Gibbs free energy 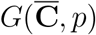, where *p* is the thermodynamic pressure (see Section 4.1 for additional discussion on the choice of this formulation), and can be decoupled into isochoric (*G*_iso_) and volumetric components (*G*_vol_) as [85, 93],

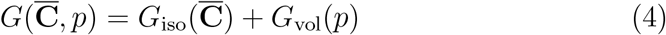

In the absence of body forces, the equations of motion are then given as,

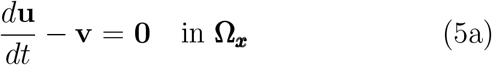

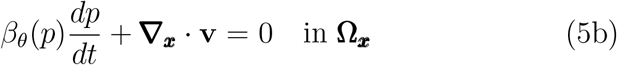

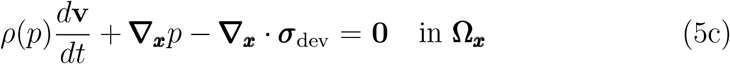

where, *β*_*θ*_ is the isothermal compressibility coefficient, *ρ* is the density of the material, ***σ***_dev_ represents the deviatoric part of the Cauchy stress tensor, and **∇**_***x***_ is the gradient operator defined in the spatial coordinates. In the above formulation, *p* is an independent variable while *ρ* and *β*_*θ*_ depend on *p*. Further, the first equation (Eq. (5a)) represents the kinematic relation between the displacement and the velocity of the body, and the following two equations (Eqs. (5b, 5c)) represent the conservation of mass and linear momentum, respectively. If *ρ*_0_ is the density of the material in the reference configuration, the constitutive relations of the hyperelastic material are represented in terms of the specific Gibbs free energy components (per unit mass) as [85, 93],

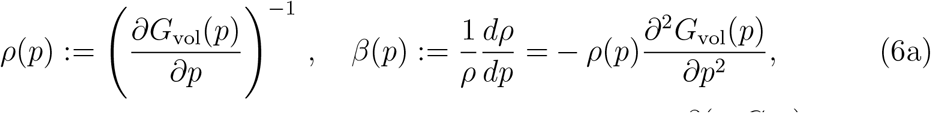

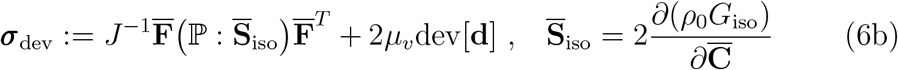

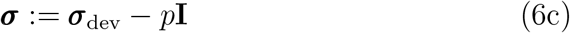

where, *µ*_*v*_ is the dynamic shear viscosity, dev[**d**] is the deviatoric part of the rate of deformation tensor, 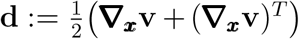,and 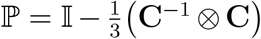 is the projection tensor. The first term in Eq. (6b) for ***σ***_dev_ represents the isochoric stress, comprising passive elastic and active contraction components, while the second term is the viscous shear stress.

We apply physiologically relevant boundary conditions on the LA myocardium. Hemodynamic pressure (*p*_la_) is applied on the LA endocardium (**Γ**_endo_), while mixed Robin-type boundary conditions are applied on the epicardium (**Γ**_epi_) and the annular cross-sections of LPV (**Γ**_lpv_), RPV (**Γ**_rpv_), and mitral valve (**Γ**_mv_), as [32, 85],

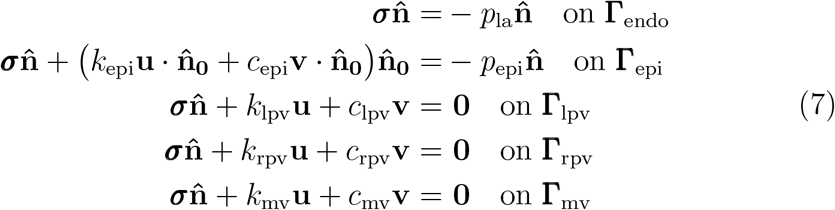

where *k*_(•)_ and *c*_(•)_ are the stiffness and damping coefficients of the Robin boundaries, respectively, and *p*_epi_ is the thoracic cavity pressure acting on the epicardium, which is typically small and is set to zero in this work, and 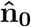 is the unit surface normal in the reference configuration. In Eq. (7), the pressures, *p*_la_ and *p*_epi_, act as follower pressure loads (i.e., the magnitude of the load changes with deformation proportional to the deformed surface area). Further, the linear stiffness and damping effects on Robin boundaries act only along the surface normal for the epicardium but act along all directions for the annular surfaces (LPV, RPV, and MV).

The equations (Eqs. (5–7)) complete the description of the initial-boundary value problem for the LA myocardial mechanics.

#### Constitutive models

Here, we describe the constitutive models used in Eq. (6) to represent the isochoric (*G*_iso_) and volumetric (*G*_vol_) free energy potentials. The isochoric myocardial deformation is split into two components – a passive elastic component and an active contraction component. We model the passive elastic response of the myocardium assuming a hyperelastic, nearly incompressible material, using the structurally-based, orthotropic Holzapfel-Ogden (HO) model in its decoupled form [94]. On the other hand, the LA contraction is modeled using the active stress-based approach. These are mathematically expressed as,

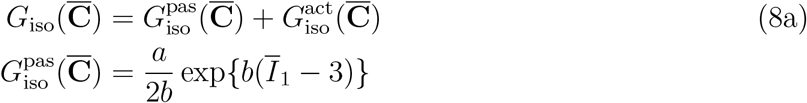

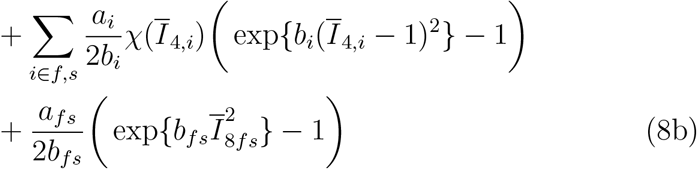

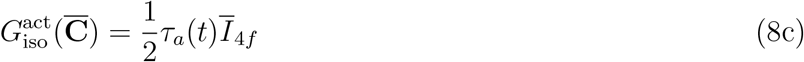

where 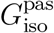 is the passive part of the isochoric Gibbs free energy, and 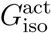 governs the active contraction. The set {*a, b, a*_*f*_, *b*_*f*_, *a*_*s*_, *b*_*s*_, *a*_*fs*_, *b*_*fs*_} defines the parameters for the HO model. Further, 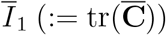 is the isotropic invariant, 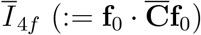 and 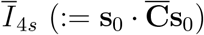 are the transverse invariants for the fiber and sheet directions, respectively, and 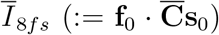 is the anisotropic invariant that captures the fiber-sheet interactions. The decoupled deformation gradient tensor 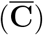 is related to the right Cauchy-Green deformation gradient tensor (**C**) using Eq. (3). We also incorporate a smooth approximation of the Heaviside function, 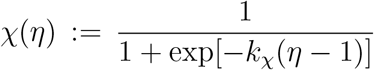, to avoid any numerical instabilities under small compressive strains [32, 95].

The additively split isochoric free energy (Eq. (8a)) translates to an additively split isochoric stress 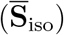 into passive 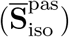 and active stress 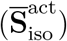 components. While the passive stress component is determined by the usual differentiation rule (Eq. (6b)) applied to the passive free energy function, 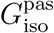 (Eq. (8b)), in this work, we adopted a phenomenological model for active stress 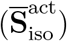,governed by a time-dependent active stress variable, *τ*_*a*_(*t*), acting along the myofibers **f**_0_ defined in the reference configuration. These are mathematically expressed as [32, 86, 95],

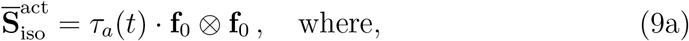

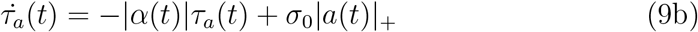

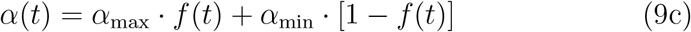

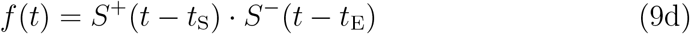

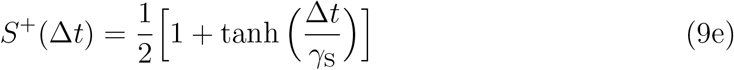

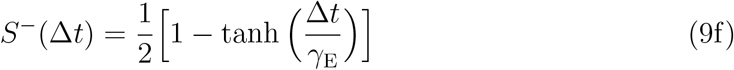

with activation function *α*(*t*), contractility *σ*_0_, and maximum and minimum activation rates, *α*_max_ and *α*_min_, respectively. Further, *S*^+^ and *S*^−^ are the descending and ascending sigmoid functions, *t*_S_ and *t*_E_ are the start and end times of contraction, and *γ*_S_ and *γ*_E_ control the steepness of contraction and relaxation, respectively. Unlike previous works [32, 95], here we used different coefficients to control the steepness of the active stress profile during contraction (*γ*_S_) and relaxation (*γ*_E_) to improve the agreement with image-based cavity volumes (Fig. 8).

To model the near-incompressibility of the myocardium, we employ the ST91 volumetric strain energy model developed by Simo and Taylor in 1991 [96], given in the form of specific Gibbs free energy as [93],

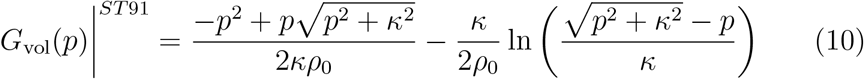

where *κ* is the bulk modulus and *ρ*_0_ is the density in the reference configuration. The corresponding *ρ*(*p*) and *β*_*θ*_(*p*) (Eq. 6a) are given by,

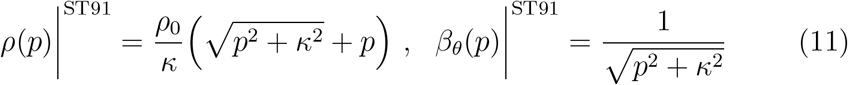

#### FE discretization, numerical methods, and solver parameters

The FEA-ready LA myocardium (Section 2.2.1, Fig. 1A) is meshed with four-noded tetrahedral (TET4) elements using Tetgen^¶^, accessible within SimVascular^‖^. A mesh convergence study was undertaken to obtain an optimal mesh size that yielded reasonable accuracy compared to the finest grid (Section 4.2). Our choice of TET4 elements was motivated by their versatility and robustness in meshing complex shapes, such as patient-specific LA myocardial geometries derived from CTA images. However, resolving finite, incompressible deformations with linear tetrahedra often leads to volumetric locking [97]. To overcome these locking issues, we adopted the variational multiscale (VMS) formulation to spatially discretize and solve the governing equations (Eqs. (5–7)). The VMS formulation allows one to employ equal-order interpolation for velocity and pressure basis functions (P1-P1 for TET4), circumventing the *inf-sup* conditions while mitigating volumetric locking, particularly in the near-incompressible limit [85, 93, 98].

Simulations are performed using an in-house multiphysics finite element solver, adapted from the open-source finite element solver, *svFSI* ^**^ [99], which was verified for cardiac mechanics applications [85, 95], and validated and employed for other cardiovascular biomechanics applications, including simulating cardiac electrophysiology [10], multiscale myocardial mechanics [86], blood flow in coronaries [100, 101], developing ventricles [22, 102, 103], fluid-structure interaction modeling in aortic dissection and aneurysms [104, 105, 106, 107], and multiphysics modeling for pediatric applications [108] and vascular growth and remodeling [109, 110]. The nonlinear system of equations is solved using the Newton-Raphson method, embedded within a predictor-multi-corrector algorithm, and integrated in time using the implicit generalized-*α* method [85, 93, 98, 111]. Within each nonlinear iteration, a block matrix equation of the type **Ax** = **B** is solved. In all the simulations performed in this study, we employed the iterative solver, generalized minimal residual (GMRES), to solve the sparse system of linear equations [112]. For the coupled 0D/3D multiscale mechanics simulations, a resistance-based preconditioning is employed to accelerate the convergence of the GMRES linear solver [86, 113]. However, for the 3D passive mechanics simulations during iFEA, the linear solver convergence is accelerated using a thresholded incomplete LU (ILUT) preconditioner, included in the Trilinos linear solver library^††^. For all the simulations, we set the tolerances for the nonlinear and the linear solvers to 10^−6^. We further set the spectral radius of infinite time step (*ρ*_**∞**_) in the generalized-*α* method to 0.5, which guarantees second-order accuracy while ensuring optimal dissipation of high-frequency temporal oscillations [111].

#### 2.2.3. Coupled 3D/0D multiscale model of LA mechanics

The LA multiscale mechanics model couples the 3D myocardial mechanics model with a 0D LPN model of the blood circulatory system [86, 87]. The 0D LPN model comprises segregated blocks for the right atrium (RA), right ventricle (RV), left ventricle (LV), and the systemic and pulmonary arterial and venous circulatory systems, connected in a closed loop. The pressure-volume relationship of the heart chambers (RA, RV, LV) is modeled using the time-varying elastance function, employing a constant passive stiffness and a double-Hill activation model for contraction (Eq. (A.5) in Appendix A) [114]. The heart valves are modeled as resistive diodes, while the systemic and pulmonary circulation blocks are modeled using resistor-inductor-capacitor (R-L-C) circuit connectors [86, 87]. The 0D model is governed by a set of differential-algebraic equations (DAEs), described in Appendix A.

The two subproblems (i.e., 3D mechanics and 0D LPN) are modularly coupled using a robust implicit coupling strategy [86, 115]. Briefly, in our coupling scheme, the rate of change of LA volume (*Q*_*LA*_) is passed from the 3D problem to the 0D model, while the LPN-computed LA pressure (*P*_*LA*_) is applied as a Neumann boundary condition on the LA endocardial surface (Fig. 4). This exchange between the 3D and the 0D solvers is done for every nonlinear (Newton) iteration until both solvers converge in a given time step. We used an implicit-explicit (IMEX) time integration strategy, where the nonlinear mechanics equations (Eqs. (5a-5c)) are integrated using the generalized-*α* method, while the coupled DAEs of the 0D LPN model are integrated using a 4^*th*^-order Runge-Kutta (RK4) time integration method.

### 2.3. Multi-step personalization procedure

The major steps involved in personalizing the multiscale mechanics model are (i) estimating the 0D LPN model parameters, (ii) characterizing the passive mechanics, i.e., estimating the HO constitutive model parameters (Eq. (8b)) and the reference stress-free configuration, and (iii) estimating the contraction model parameters (Eq. (9)) such that the FEA-predicted deformation aligns with image data over the entire cycle. These steps are described in the following subsections.

#### 2.3.1. 0D LPN parameter estimation

To personalize the 0D LPN component of the multiscale mechanics model (Fig. 4), the 3D FE model is replaced with a 0D surrogate model of LA, resulting in a fully 0D model of blood circulation (Fig. 5, left). Similar to the other cardiac chambers, the pressure-volume relationship of this surrogate LA is modeled using a time-varying elastance function, characterized by a constant passive stiffness and a double-Hill activation function to model contraction (Eq. (A.5) in Appendix A) [114]. The following optimization procedure is then performed to estimate the LPN parameters that match the patient data.

A set of hemodynamic pressure measurements and image-based volumes is defined as ‘targets’ for the optimization method based on the available clinical data for the patient and from prior studies (Table 2). Target quantities based on the patient’s clinical data include (i) minimum and maximum cardiac volumes 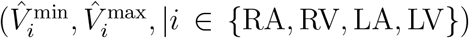 extracted from the CTA images by applying a deep-learning based automatic segmentation algorithm, HeartDeformNet [116], and (ii) diastolic (minimum) and systolic (maximum) cuff-based pressures 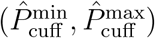.To enhance the robustness of the optimization and yield physiologically relevant results, we augmented the clinically available data with literature-based pressure data, such as the minimum and maximum pulmonary artery pressures 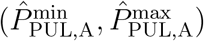 [117], pulmonary artery wedge pressure 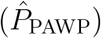 [117, 118], and central venous pressure 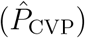 [117, 119, 120]. All these targets are summarized in Table 2.

**Table 2:**
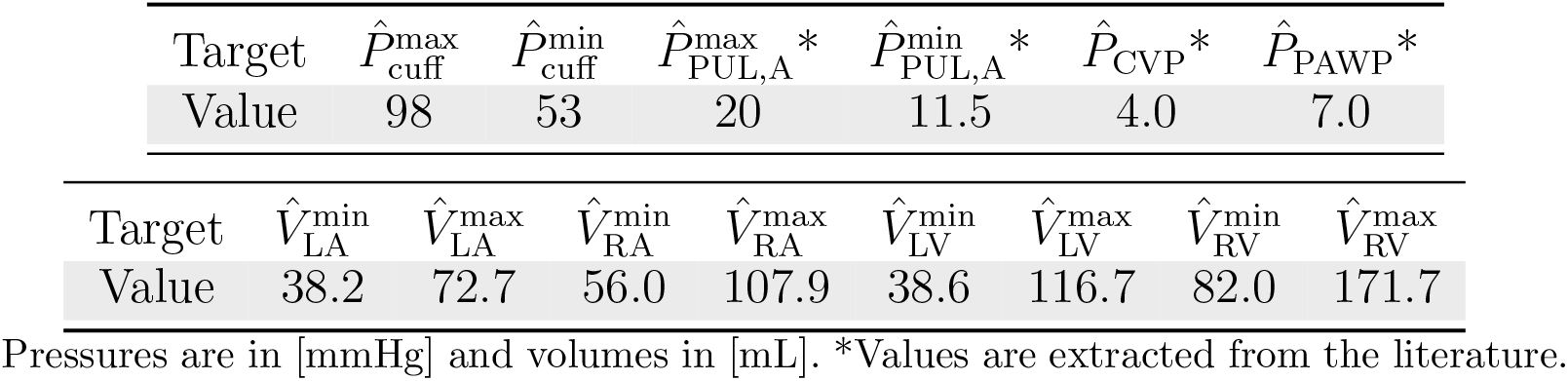
Clinically available patient data and literature-based pressure values are set as ‘target’ values for the LPN parameter estimation framework. LA: left atrium, RA: right atrium, LV: left ventricle, RV: right ventricle, PUL,A: pulmonary artery, CVP: central venous pressure, PAWP: pulmonary artery wedge pressure.

All the LPN parameters are initialized with values from previous studies [87, 121]. The patient’s ECG data is used to compute the activation start and end times of the time-varying elastance function for each cardiac chamber, as discussed in Appendix B. Further, only the parameters proximal to the left heart and the most sensitive ones, based on the total Sobol index at least 0.2 [9], are tuned as part of the optimization. The remaining parameters are fixed to accelerate convergence and reduce computational cost. The final sets of tunable LPN parameters are,

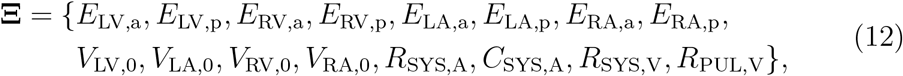

comprising the active (*E*_*i*,*a*_) and passive (*E*_*i*,*p*_) stiffness parameters of the cardiac chambers (*i* **∈** {LV, RV, LA, RA}), the resting volumes of LA (*V*_LA,0_), LV (*V*_LV,0_), RA (*V*_RA,0_), and RV (*V*_RV,0_), the systemic arterial resistance (*R*_SYS,A_) and compliance (*C*_SYS,A_), the systemic venous resistance (*R*_SYS,V_), and the pulmonary venous resistance (*R*_PUL,V_).

In our optimization scheme, only deviations above a certain threshold (*ϵ*_*θ*_) are penalized. The threshold is chosen as 5% for the clinically measured targets but 20% for quantities derived from the literature (Table 2). The objective function is thus defined as,

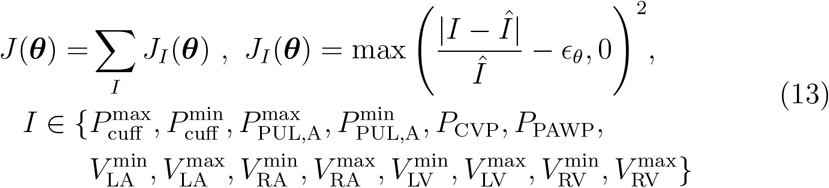

where *Î* and *I* are the optimization target quantities (Table 2) and the corresponding simulated values, respectively. We used the box-constrained, limited Broyden-Fletcher-Goldfarb-Shanno (`L-BFGS-B`) algorithm in the `scipy.optimize` library to minimize the above objective function (Eq. (13)). Simulations are performed until a limit cycle is reached, typically after 30 cardiac cycles, before comparing against the clinical targets. The tunable parameters are bounded within 25% of its initial value, i.e., Ξ_*i*_ **∈** [0.75Ξ_*i*,0_, 1.25Ξ_*i*,0_], where Ξ_*i*,0_ is the initial parameter value. Lastly, the optimization is repeated for ten random parameter initializations within the bounds, and the best-fit values are used for subsequent analysis.

#### 2.3.2. Personalizing passive mechanics

After optimizing the surrogate-based LPN model (Section 2.3.1), the LA pressure profile during passive expansion is used to personalize the 3D myocardial passive mechanics. This idea to predict myocardial passive material parameters to match the image-based inner boundary deformations was first proposed by Ghista et al. [122] as mechano-myocardiography (MMG) for left ventricular myocardium, followed by several variants in later studies [123, 124, 125, 126, 127, 128, 129, 130, 131, 132]. Here, we adopted our recently developed inverse finite element analysis (iFEA) framework that involves a multi-level optimization scheme to extract passive myocardial properties and simultaneously estimate the unloaded reference configuration to match deformations from gated CTA [85]. The iFEA algorithm includes an outer level iterating on the material parameters to match image-based LA volumes and local nodal displacements at select landmarks (Fig. 5), while an inner level estimates the unloaded reference configuration using Sellier’s algorithm for a combination of material parameters and pressure load set by the outer level. This nested optimization process enables extracting patient-specific LA tissue parameters that fit time-dependent, image-based LA topology under personalized pressure loads. The method is applied for estimating the six HO constitutive model parameters ( {*a, b, a*_*f*_, *b*_*f*_, *a*_*s*_, *b*_*f*_}, Eq. (8b)) to fit LA volumes and displacements at three landmarks during passive expansion (20–40% R-R interval) [85]. While the current work employs a custom-designed genetic algorithm for the outer-level iterations due to its superior performance, the reader is referred to Shi et al. [85] for additional technical details on the choice of constitutive model parameters for tuning, algorithm implementation, optimization methods, and other sensitivity analyses.

#### 2.3.3. Active contraction tuning

We then focus on estimating the contraction model parameters to personalize LA deformation over the entire cycle. Here, we used the multiscale mechanics model coupling the 0D LPN with the 3D FE model of LA (Fig. 4). The LPN parameters are extracted from the optimized full 0D model (Section 2.3.1) while the 3D unloaded reference configuration and passive myocardial constitutive parameters are retrieved from the iFEA analysis (Section 2.3.2). We then iteratively tuned the active stress model parameters, *σ*_0_ and *γ*_E_ (Eq. (9)), until the LA chamber volumes and deformation closely aligned with the data from CTA images. The activation start and end times (*t*_S_, *t*_E_) times are determined from ECG measurements (Appendix B), while the remaining parameters (*α*_max_, *α*_min_, *γ*_S_) are extracted from literature [32], and are kept unchanged.

### 2.4. Moving-domain blood flow formulation

Once the multiscale mechanics simulations converge to a periodic solution (Section 2.3), the deformation of the inner wall is extracted from the last cycle and imposed as a moving-wall boundary condition to drive blood flow through the LA cavity. We modeled blood flow as incompressible and Newtonian, and its motion in moving domains is governed by the arbitrary Lagrangian-Eulerian (ALE) formulation of the Navier-Stokes equations, given by [22],

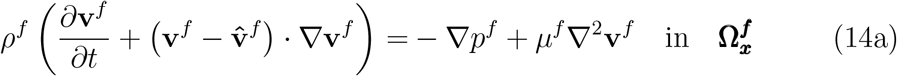

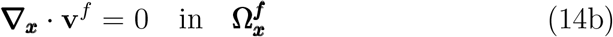

where,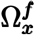 is the fluid domain comprising the LA blood pool (Fig. 6A), extracted as the capped inner wall (endocardium) from the updated myocardial configuration at the beginning of the last cycle from the multiscale mechanics simulation. 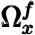 is bounded by 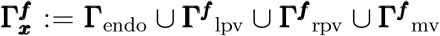,such that **Γ**_endo_ is the LA endocardium (inner wall), whereas **Γ**^***f***^ _lpv_, **Γ**^***f***^ _rpv_, and **Γ**^***f***^ _mv_ represent the capped planes of LPV, RPV, and MV, respectively, through which the blood flows into and out of the LA cavity. These capping planes are created using SimVascular’s ‘fill hole’ feature so that the resulting fluid domain is watertight. *ρ*^*f*^ and *µ*^*f*^ are the density and dynamic viscosity of blood, **v**^*f*^ and *p*^*f*^ are the local fluid velocity and pressure, and 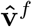 is the velocity of the fluid domain that is smoothly deformed to conform to the imposed endocardial motion. Eq. (14a) represents the balance of linear momentum for an incompressible Newtonian flow in ALE coordinates, while Eq. (14b) represents the continuity equation for mass balance. To close the above equations, the boundary conditions are applied as,

**Figure 6:**
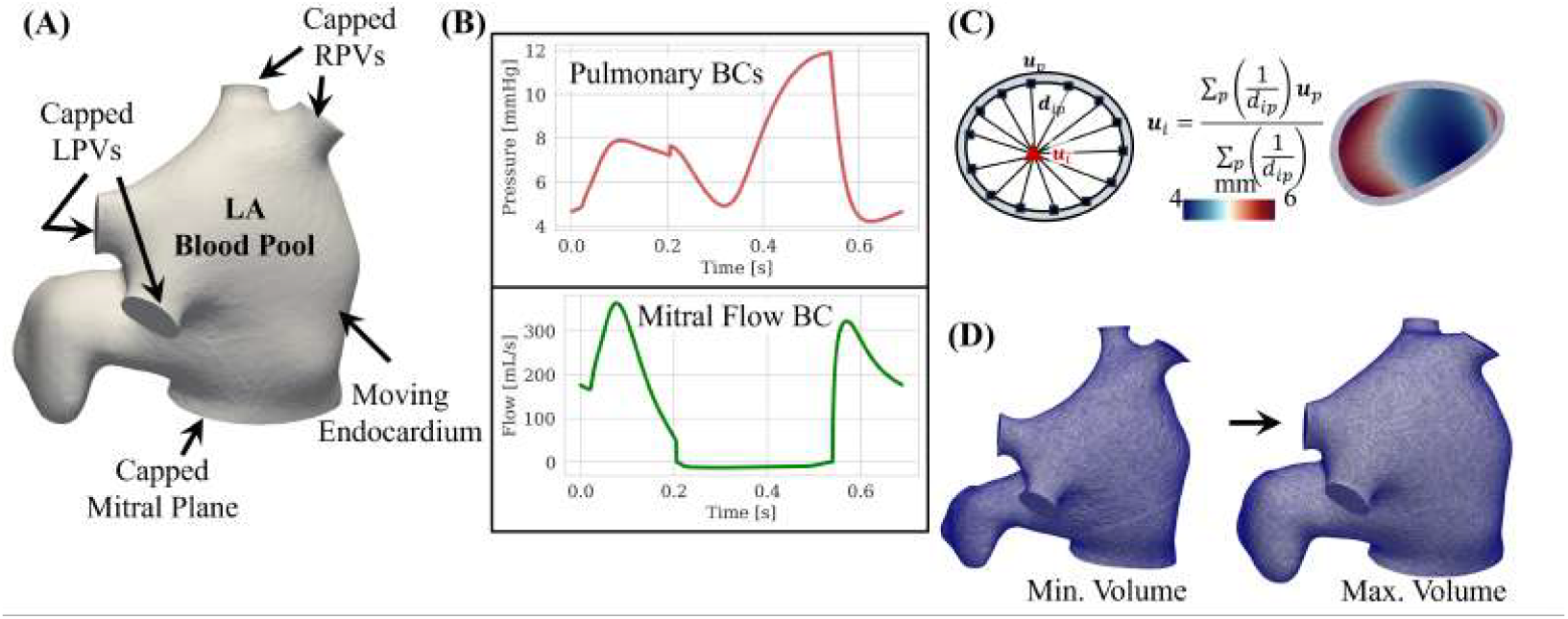
Moving domain blood flow simulation set up using data from personalized multiscale mechanics modeling of the left atrium (LA). (A) LA blood pool constructed from the inner wall at the beginning of the last cycle of multiscale mechanics simulations, with the pulmonary veins and mitral plane capped using the ‘fill hole’ feature in SimVascular. (B) Time-dependent boundary conditions, extracted from multiscale mechanics simulations, are imposed on the pulmonary veins (top) and the mitral plane (bottom). (C) An inverse distance weighted (IDW)-based projection method is adopted to smoothly project the tissue displacements at the perimeter of the capped boundaries to the interior nodes. (D) Fluid mesh representations at the minimum and maximum volumes during the cycle.

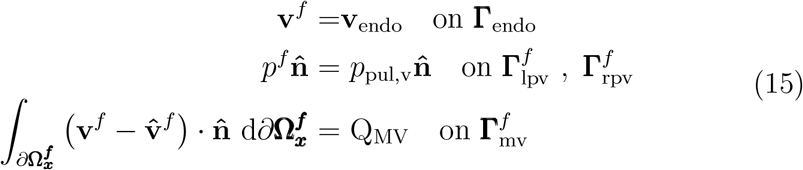

where, **v**_endo_ is the endocardial velocity, *p*_pul,v_ is the pressure at the pulmonary veins (Fig. 6B, top), and Q_MV_ is the flowrate through the mitral plane (Fig. 6B, bottom). All these time-dependent quantities are extracted from the multiscale myocardial mechanics simulation (Section 2.3, Fig.4). In particular, *p*_pul,v_, computed as (*P*_PUL,V_ − *Q*_PUL,V_*R*_PUL,V_), is assumed to be the same on LPV and RPV boundaries 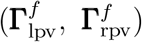, and a uniform velocity profile is assumed at the mitral plane for prescribing the atrioventricular flux, Q_MV_ [41, 133, 134].

Further, as part of the ALE formulation, the fluid domain is smoothly deformed by solving an element-Jacobian-stiffened linear elastostatics problem for the fluid mesh velocity 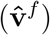,augmented by Dirichlet boundary conditions at the LA endocardium (i.e.,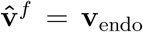 on **Γ**_endo_) and the inflow/outflow planes 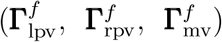. However, as the velocities of these capped inflow/outflow planes are not explicitly available from the myocardial mechanics simulations, we employ an inverse distance weighting (IDW)-based projection method to interpolate the local nodal velocities of the capped planes from the tissue velocity at the corresponding perimeters. The weights are computed using an inverse-distance measure between an interior node on the capped surface and each perimeter node (Fig. 6C). This results in a smooth velocity projection from the cap perimeter to the cap interior, which is repeated throughout the cardiac cycle. Ultimately, our setup will result in a fluid-tight LA cavity that smoothly deforms due to the motion imposed on the inner wall extracted from the multiscale mechanics simulations and drives the blood flow under personalized boundary conditions at the pulmonary veins and the mitral plane (Fig. 6D).

#### 2.4.1. FE discretization, numerical methods, and solver parameters

Similar to the LA myocardium, the blood pool or lumen is meshed with TET4 elements using Tetgen^*^, accessible within SimVascular^†^. To account for the strong advection, we employed a finer mesh resolution for blood flow analysis, resulting in about 1.9M TET4 elements (Fig. 6D). VMS stabilization is employed to circumvent *inf-sup* conditions for incompressibility with mixed finite elements and address convection-related instabilities [22, 93, 135], thereby allowing us to use equal-order interpolations for velocity and pressure basis functions (P1-P1 for TET4). The system of equations (ALE-based fluid flow and mesh motion) is solved using the same in-house parallelized multiphysics finite element solver employed for multiscale myocardial mechanics modeling. The equations are integrated in time using the generalized-*α* method, and the nonlinearity is handled using a block Newton method, embedded within a predictor-multi-corrector algorithm [93, 135].

Here, for each Newton iteration, the ALE system of equations (Eq. 14) and the mesh motion equations are solved in a loosely coupled manner. Further, within each block-wise nonlinear iteration, we employed the GMRES linear solver with diagonal preconditioning to solve the linearized ALE system of equations, whereas the Conjugate Gradient (CG) method is used to solve the linearized system of equations governing the fluid mesh motion.

All the tolerances (linear and nonlinear) are set to 1e-6 for convergence, while the spectral radius of infinite time-step (*ρ*_**∞**_) is set to 0.5. We choose 0.69 ms as the time step size, resulting in approximately 1000 timesteps per cardiac cycle. A linear interpolation in time is applied to compute the imposed nodal boundary velocities at intermediate time steps, whereas a Fourier interpolation is applied for the pulmonary pressures and mitral flow rates.

The simulations are run for three cycles to remove the initial transient and arrive at a quasi-periodic state. In the current simulations, we did not apply any dynamic remeshing to handle large deformations. The simulations are run using 384 AMD-EPYC-7742 CPU cores on the Expanse supercomputing cluster at UC San Diego, allocated through the NSF-sponsored ACCESS program, taking approximately 9.5 hours for one cardiac cycle.

## 3. Result

### 3.1. Performance of 0D LPN parameter estimation and iFEA

Parameters obtained from optimizing the fully 0D LPN model, with a surrogate model for LA (Section 2.3.1), are shown in Table 3. A comparison of the optimized LPN-predicted hemodynamic pressures and chamber volumes shows a reasonable agreement with the clinical data (Fig. 7A). While the relative error in most of these quantities is within 5%, the relative errors in min/max LA volume 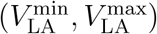 and central venous pressure (*P*_CVP_) are below 10%. The minimum pulmonary artery pressure 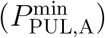 and pulmonary artery wedge pressure (*P*_PAWP_) have the highest relative errors of 22% and 31%, respectively, although the absolute differences are only a couple of millimeters of Hg, attributed to the lack of actual patient data and high uncertainty in the literature values [117, 118, 119, 120]. The P–V loop from this simplified 0D surrogate model of LA mechanics captures the *a*–loop but fails to capture the *v* –loop (Fig. 7B). All other hemodynamic indices from the 0D LPN are obtained within reasonable physiological limits (Fig. 9).

**Table 3:** Optimized parameters for the fully 0D LPN model. Only the most sensitive parameters (marked by *), with a total Sobol index of at least 0.2 [9], are varied during optimization. Reference values are used for all other parameters [87, 121]. The parameters and the differential-algebraic equations (DAEs) governing the LPN model are described in Appendix A.

| *Cardiac parameters* |  |  |  |
| --- | --- | --- | --- |
| LV active elastance* | $E_{LV,a}$ | 12.5 | mmHg/mL |
| LV passive elastance* | $E_{LV,p}$ | 0.075 | mmHg/mL |
| RV active elastance* | $E_{RV,a}$ | 1.53 | mmHg/mL |
| RV passive elastance* | $E_{RV,p}$ | 0.0375 | mmHg/mL |
| LA active elastance* | $E_{LA,a}$ | 0.210 | mmHg/mL |
| LA passive elastance* | $E_{LA,p}$ | 0.259 | mmHg/mL |
| RA active elastance* | $E_{RA,a}$ | 0.150 | mmHg/mL |
| RA passive elastance* | $E_{RA,p}$ | 0.075 | mmHg/mL |
| LV rest volume* | $V_{LV,0}$ | 31.4 | mL |
| RV rest volume* | $V_{RV,0}$ | 73.2 | mL |
| LA resting volume* | $V_{LA,0}$ | 30.3 | mL |
| RA resting volume* | $V_{RA,0}$ | 41.5 | mL |
| *Circulation parameters* |  |  |  |
| Systematic arterial resistance* | $R_{SYS,A}$ | 0.420 | mmHg · s/mL |
| Systematic arterial compliance* | $C_{SYS,A}$ | 1.403 | mL/mmHg |
| Systematic aortic inductance | $L_{SYS,A}$ | 0.005 | mmHg · s<sup>2</sup>/mL |
| Systematic venous resistance* | $R_{SYS,V}$ | 0.228 | mmHg · s/mL |
| Systematic venous compliance | $C_{SYS,V}$ | 60.0 | mL/mmHg |
| Systematic venous inductance | $L_{SYS,V}$ | 0.0005 | mmHg · s<sup>2</sup>/mL |
| Pulmonary arterial resistance | $R_{PUL,A}$ | 0.032 | mmHg · s/mL |
| Pulmonary arterial compliance | $C_{PUL,A}$ | 10.0 | mL/mmHg |
| Pulmonary arterial inductance | $L_{PUL,A}$ | 0.0005 | mmHg · s<sup>2</sup>/mL |
| Pulmonary venous resistance* | $R_{PUL,V}$ | 0.035 | mmHg · s/mL |
| Pulmonary venous compliance | $C_{PUL,V}$ | 16.0 | mL/mmHg |
| Pulmonary venous inductance | $L_{PUL,V}$ | 0.0005 | mmHg · s<sup>2</sup>/mL |
| *Valve parameters* |  |  |  |
| Valve opening resistance | $R_{min}$ | 0.005 | mmHg · s/mL |
| Valve closing resistance | $R_{max}$ | 5.0 | mmHg · s/mL |

**Figure 7:**
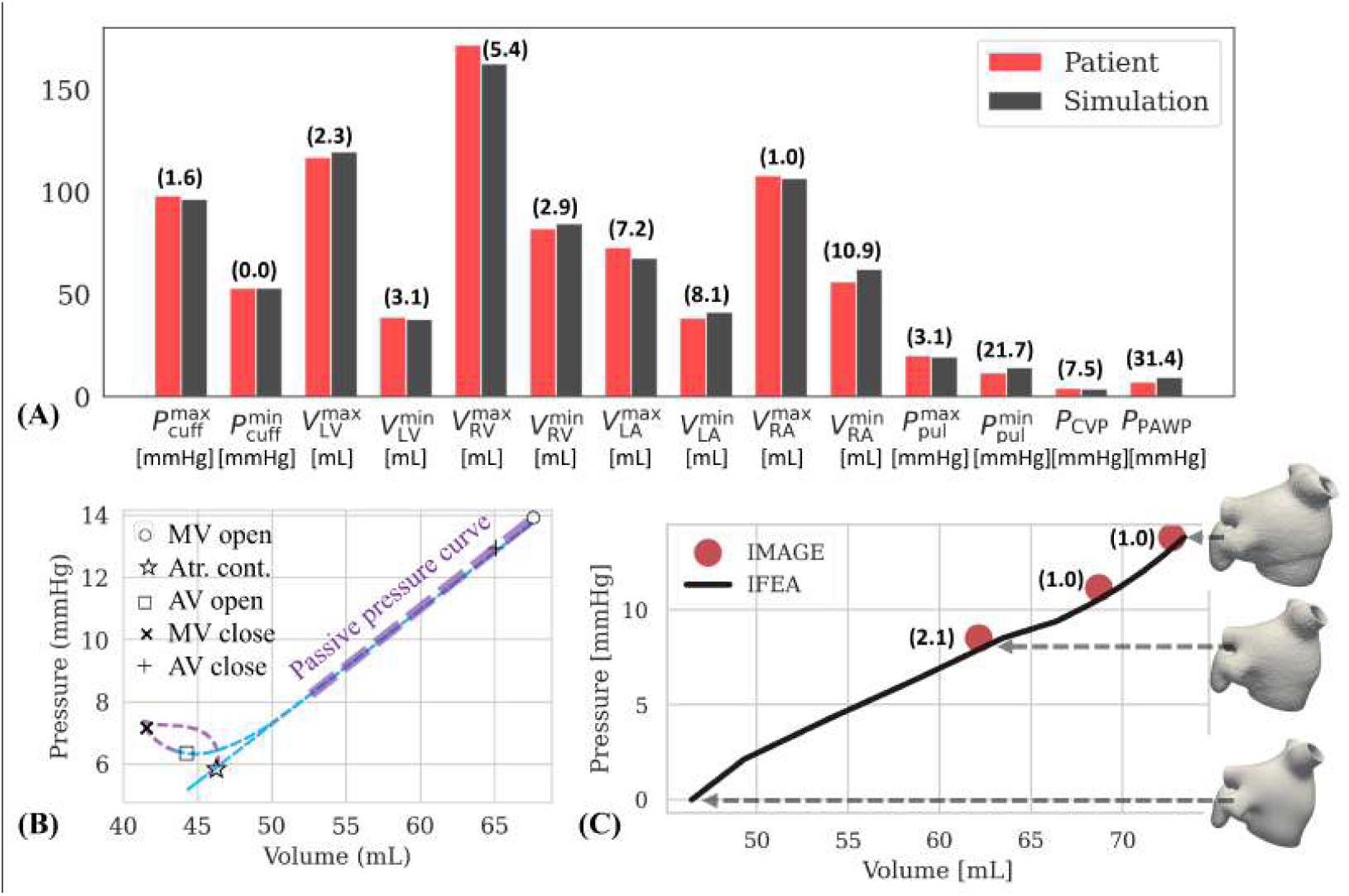
(A) Comparison of optimized LPN-predicted pressures and cardiac chamber volumes against clinical data, with % relative errors in parentheses. (B) LPN-predicted P–V loop, highlighting the pressure profile during passive expansion (20-40% R-R) and the opening and closing states of the mitral valve (MV) and aortic valve (AV). (C) Estimated reference (unloaded) configuration from iFEA. Passive loading with iFEA-optimized material parameters and LPN-based pressure profile shows close agreement between simulated LA volumes and image data. Values in parentheses indicate % relative errors. LA/RA: left/right atrium; LV/RV: left/right ventricle; PAWP: pulmonary artery wedge pressure; CVP: central venous pressure; LPN: lumped parameter network; P–V: pressure–volume; iFEA: inverse finite element analysis.

**Figure 8:**
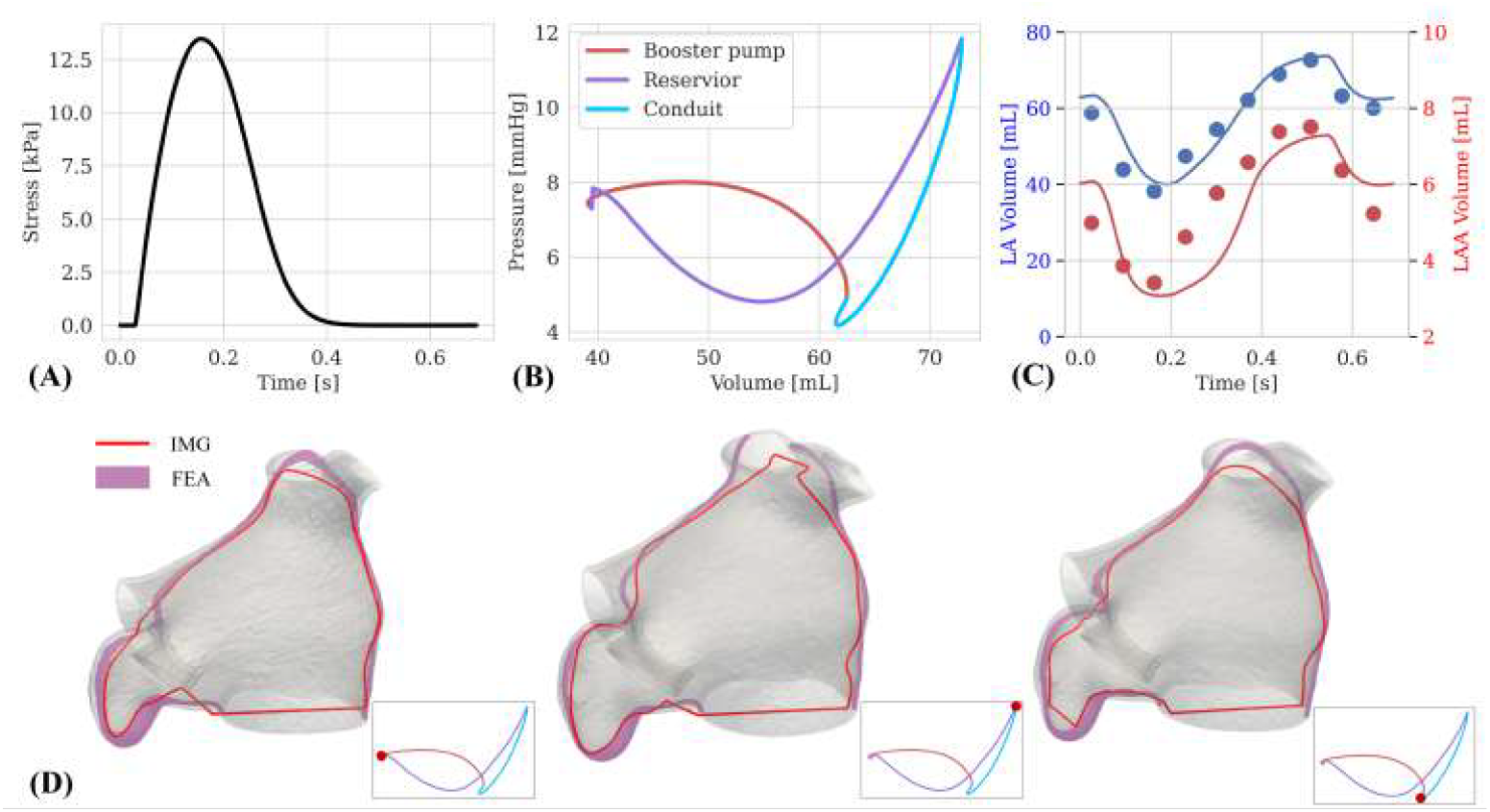
(A) Time-varying active stress during a cardiac cycle with optimized parameters (Table 5). (B) LA pressure–volume (P–V) diagram from fully personalized multiscale mechanics simulations, capturing both *a*– and *v* – loops, comprising conduit, reservoir, and booster pump function. (C) Comparison of time-varying LA and LAA cavity volumes from personalized multiscale mechanics simulations against image data at all phases during the cardiac cycle. (D) Comparison of LA deformation against image slices at three selected phases during the cycle - (left) end of contraction, (center) end of expansion, and (right) beginning of contraction. LA: left atrium; LAA: left atrial appendage.

**Figure 9:**
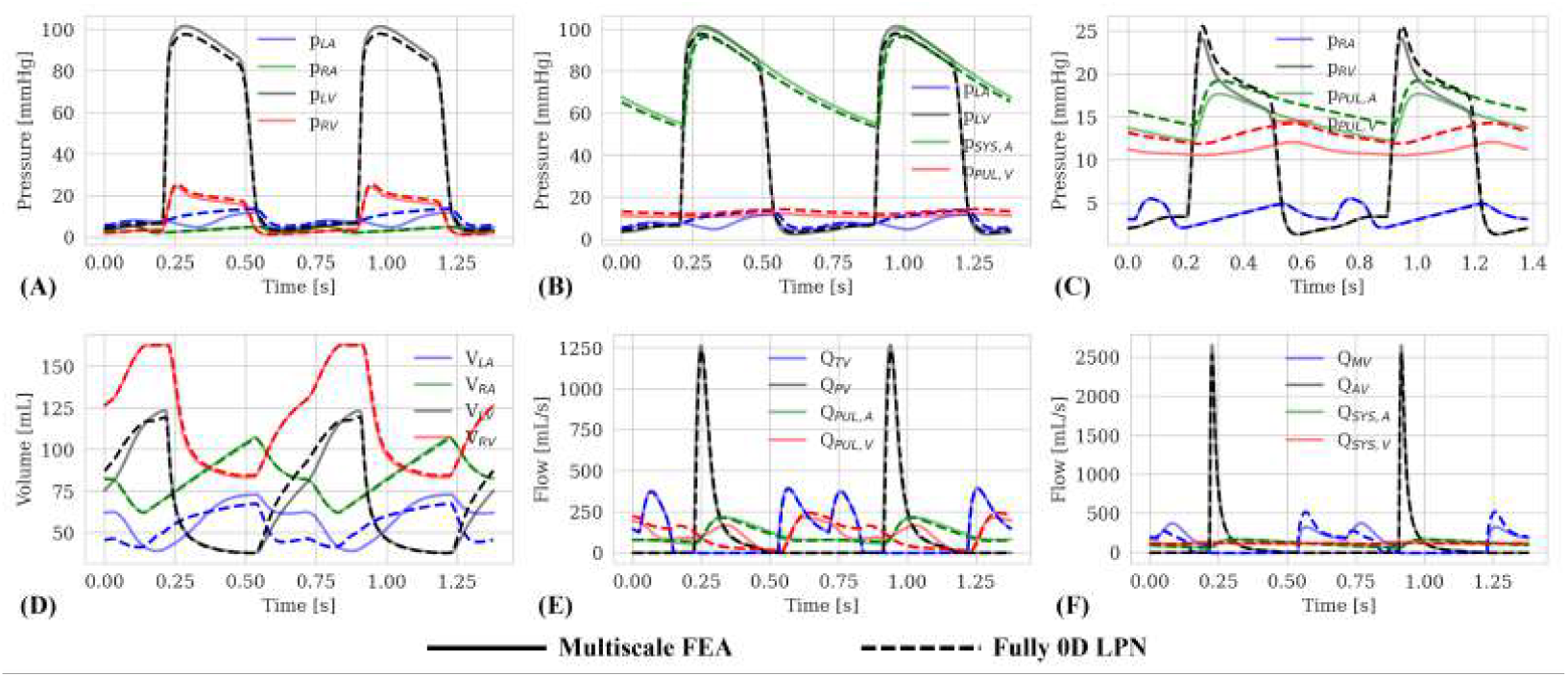
Comparison of time-dependent LPN data between coupled 0D-3D multiscale FEA simulation and fully 0D LPN with a surrogate LA model. LA: left atrium; LPN: lumped parameter network.

The passive pressure profile from the personalized 0D model (highlighted in Fig. 7B) is used as the time-varying endocardial pressure boundary condition for iFEA [85]. Simulated LA deformation using the iFEA-optimized reference configuration and material parameters (Table 4) with this passive pressure load shows a close alignment with the CTA data (Fig. 7C), with relative errors in the LA cavity volumes under 2%.

**Table 4:** Personalized parameters governing LA passive mechanics, including the optimal Holzapfel-Ogden (HO) constitutive model parameters, other myocardial material parameters, and Robin boundary condition parameters. HO model parameters, marked by *, are optimized during inverse finite element analysis (iFEA), while all other parameters remain fixed [85].

| *HO model parameters* |  |  |  |
| --- | --- | --- | --- |
| Isotropic stiffness* | $a$ | 3380 | dyn/cm<sup>2</sup> |
| Isotropic exponential coefficient* | $b$ | 5.3 | – |
| Fiber stiffness* | $a_f$ | 234870 | dyn/cm<sup>2</sup> |
| Fiber exponential coefficient* | $b_f$ | 15.5 | – |
| Sheet stiffness* | $a_s$ | 164710 | dyn/cm<sup>2</sup> |
| Sheet exponential coefficient* | $b_s$ | 11.2 | – |
| Fiber-sheet stiffness | $a_{fs}$ | 2160 | dyn/cm<sup>3</sup> |
| Fiber-sheet exponential coefficient | $b_{fs}$ | 11.4 | – |
| *Robin boundary condition parameters* |  |  |  |
| Pulmonary vein annuli stiffness | $k_{lpv}, k_{rpv}$ | 10<sup>6</sup> | dyn/cm<sup>3</sup> |
| Pulmonary vein annuli damping | $c_{lpv}, c_{rpv}$ | 500 | dyn · s/cm<sup>3</sup> |
| Epicardial stiffness | $k_{epi}$ | 2 × 10<sup>4</sup> | dyn/cm<sup>3</sup> |
| Epicardial damping | $c_{epi}$ | 500 | dyn · s/cm<sup>3</sup> |
| Mitral valve annular stiffness | $k_{mv}$ | 10<sup>4</sup> | dyn/cm<sup>3</sup> |
| Mitral valve annular damping | $c_{mv}$ | 500 | dyn · s/cm |
| *Other parameters* |  |  |  |
| Myocardium density | $\rho$ | 1.055 | g/cm<sup>3</sup> |
| Volumetric bulk modulus | $\kappa$ | 10<sup>7</sup> | dyn/cm<sup>2</sup> |
| Myocardium viscosity | $\mu_v$ | 10<sup>3</sup> | dyn · s/cm<sup>2</sup> |
| Step function parameter | $k_\chi$ | 100 | – |

Regarding computational performance, the 0D LPN parameter estimation takes about 11 hours for ten random parameter initializations on a local workstation with an Intel Core i9-13900 processor (5.8 GHz, 24 cores) and 128GB of memory. Likewise, the iFEA takes approximately six hours (30 generations) to converge using a moderate mesh size (∼ 90K tetrahedral elements) on an Intel Xeon Gold 6226 2.9GHz CPU with 32 cores.

### 3.2. Active contraction parameters and multiscale modeling

Table 5 lists the iteratively adjusted active stress model parameters to simulate LA contraction using the multiscale FEA setup (Section 2.2.3) and previously described tuning procedure (Section 2.3.3). Each multiscale simulation converges to a periodic state in about five cardiac cycles, with a total simulation time of approximately 1.5 hours using a single node on the Ginsburg high-performance computing cluster at Columbia University, with two Intel Xeon Gold 6226 CPUs (2×16 cores, 2.9GHz) and 192GB of memory.

**Table 5:** Optimized active stress model parameters to simulate LA contraction and perform multiscale FEA. *parameters varied during tuning. ^‡^parameters extracted from ECG data (Appendix B). All other parameters are extracted from literature [32].

|  |  |  |  |
| --- | --- | --- | --- |
| Minimum activation rate | $\alpha_{\min}$ | –30 | – |
| Maximum activation rate | $\alpha_{\max}$ | 5 | – |
| Contractility* | $\sigma_0$ | 360000 | dyn/cm<sup>2</sup> |
| Contraction start time† | $t_S$ | 0.025 | s |
| Contraction end time† | $t_E$ | 0.270 | s |
| Contraction steepness | $\gamma_S$ | 0.005 | – |
| Relaxation steepness* | $\gamma_E$ | 0.2 | – |

Unlike the fully 0D LPN simulation (Fig.7B), the multiscale simulations effectively capture both *a*– and *v* –loops in the P–V diagram (Fig. 8B), using the optimized active stress profile for contraction (Fig. 8A). The resulting LA deformation reasonably agrees with the image data throughout the cardiac cycle (Fig. 8C, D).

In particular, our multi-step personalization procedure for multiscale LA mechanics results in a decent agreement in the cavity volumes of LA and LAA between simulations and image data (Fig. 8C). The average relative error in LA volume change is about 6.8% while the mean error in LAA volume change is about 14%. The relative errors in the LA min/max volume, SV, and EF are 0.8%, 6.8%, 5.8%, and 6.6%, respectively, while the corresponding values for LAA are 4%, 8.8%, 0.0%, and 4.2%, respectively. Further, the overall FEA-predicted 3D LA deformation agrees qualitatively with image slices at multiple phases during the cardiac cycle (Fig. 8D).

We also compared the time-dependent LPN output between the coupled 0D-3D FEA simulation and the fully 0D LPN with a surrogate LA model (Fig. 9). Not surprisingly, most LPN variables are comparable between the two simulation setups, except for quantities directly related to the LA that show noticeable differences. These include LA pressure (*p*_LA_, Fig. 9A, B), volume (*V*_LA_, Fig. 9D), and the flowrate through the mitral valve (*Q*_MV_, Fig. 9F).

### 3.3. LA tissue mechanics using personalized multiscale FEA

Our results predict large myocardial displacements around the septum spanning the anterior and the inferior walls at the end of contraction and during the reservoir phase, when LA is at its minimum and maximum volumes, respectively (top row in Fig. 10 and Fig. C.15 in Appendix C). We also find that both LA and LAA are subjected to peak strain during contraction compared to all other phases (second row in Fig. 10 and Fig. C.15 in Appendix C), signifying its role in emptying the blood pool from its cavity during LA contraction or A-wave. However, this increased strain doesn’t translate into increased wall stress or stress concentration; instead, the LAA wall stresses appear near-uniformly distributed at all phases of the cardiac cycle (third row in Fig. 10 and Fig. C.15 in Appendix C). We also notice localized stress concentration sites around regions with sharp changes in fiber orientations, such as the region near the mid-anterior and mid-inferior walls proximal to the mitral plane.

**Figure 10:**
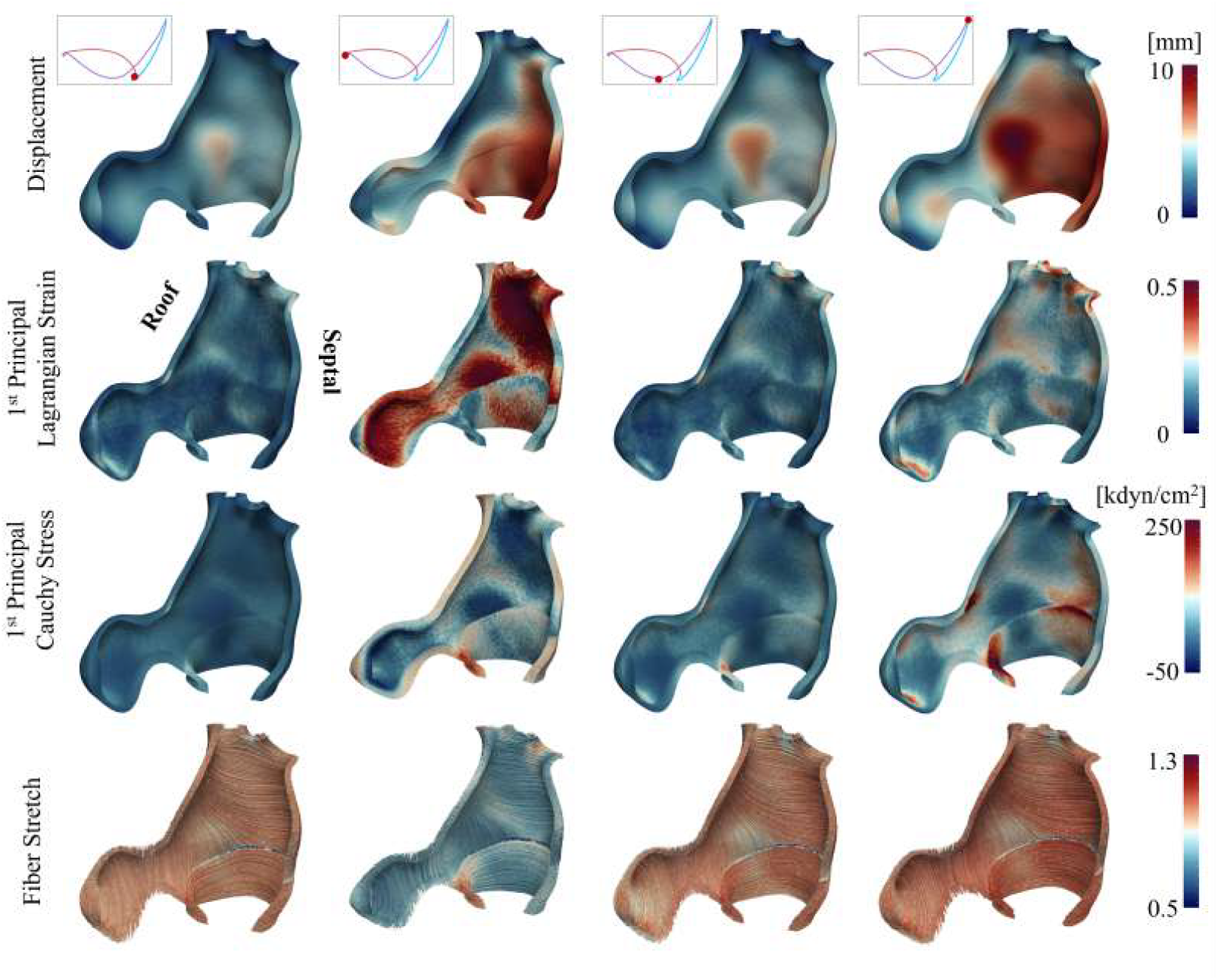
Characterizing LA mechanics along the anterior view during the cardiac cycle using the current personalized, multiscale FEA modeling framework (Fig. 2). The complementary clipped view showing the inferior wall along the posterior view is included in Appendix C. (row-wise top to bottom) displacement, 1^st^ principal (Green-) Lagrange strain, 1^st^ principal Cauchy stress, and fiber stretch. (column-wise left to right) beginning of contraction or end of conduit phase, end of contraction (booster pump), mid-expansion (reservoir), and end of expansion or beginning of the conduit phase.

As expected, the fiber stretch values are below one during contraction and exceed one during expansion (bottom row in Fig. 10 and Fig. C.15 in Appendix C). In our model, the LA fibers experience a maximum elongation of up to 30% and a peak shortening of up to 50%. However, a histogram plot suggests that fiber stretch depends on the cardiac phase (Fig. C.18 in Sec. Appendix C). During the conduit phase, the fiber stretch exhibits a near-symmetric distribution around the equilibrium state, suggesting a mix of elongation and shortening. However, during expansion, there is a rightward shift in the curve, with more elongation, although the spread suggests that the fiber stretch is non-uniform across the tissue. However, by the end of contraction, the entire curve shifts left to the other side of the spectrum, with nearly all the fibers experiencing contraction. The broader spread of the histogram during contraction indicates a greater degree of non-uniformity in the fiber stretch during contraction than expansion.

Overall, our results highlight the importance of spatial heterogeneity and cardiac phase-dependence of atrial mechanics, regulated by fiber architecture, material parameters, and boundary conditions.

### 3.4. LA blood flow using personalized and imposed wall motion

Fig. 11 shows a clipped anterior view of LA hemodynamics, including streamlines, vortex structures based on Q-criterion [136], wall shear stress (WSS), and surface pressure in a patient-specific LA model at the same four phases of the cycle as the mechanics data visualization (Fig. 10). The complementary clip depicting the corresponding quantities on the inferior wall along the posterior-anterior direction is provided in Section Appendix C. At the beginning of contraction (Fig. 11, first column) and passive expansion (Fig. 11, third column), we notice strong high-velocity jets emanating from LPVs into the atrial cavity. However, it is interesting that the jets from RPVs are relatively weaker, although they are driven by the same upstream pressure. We also notice a complex flow field with interacting vortices in the bulk of the LA cavity through the contraction and expansion phases (Fig. 11, left three columns). However, these swirling eddies decay as LA reaches its maximum volume and prepares for supplying LV with blood during the ensuing conduit and booster pump phases (Fig. 11, right-most column).

**Figure 11:**
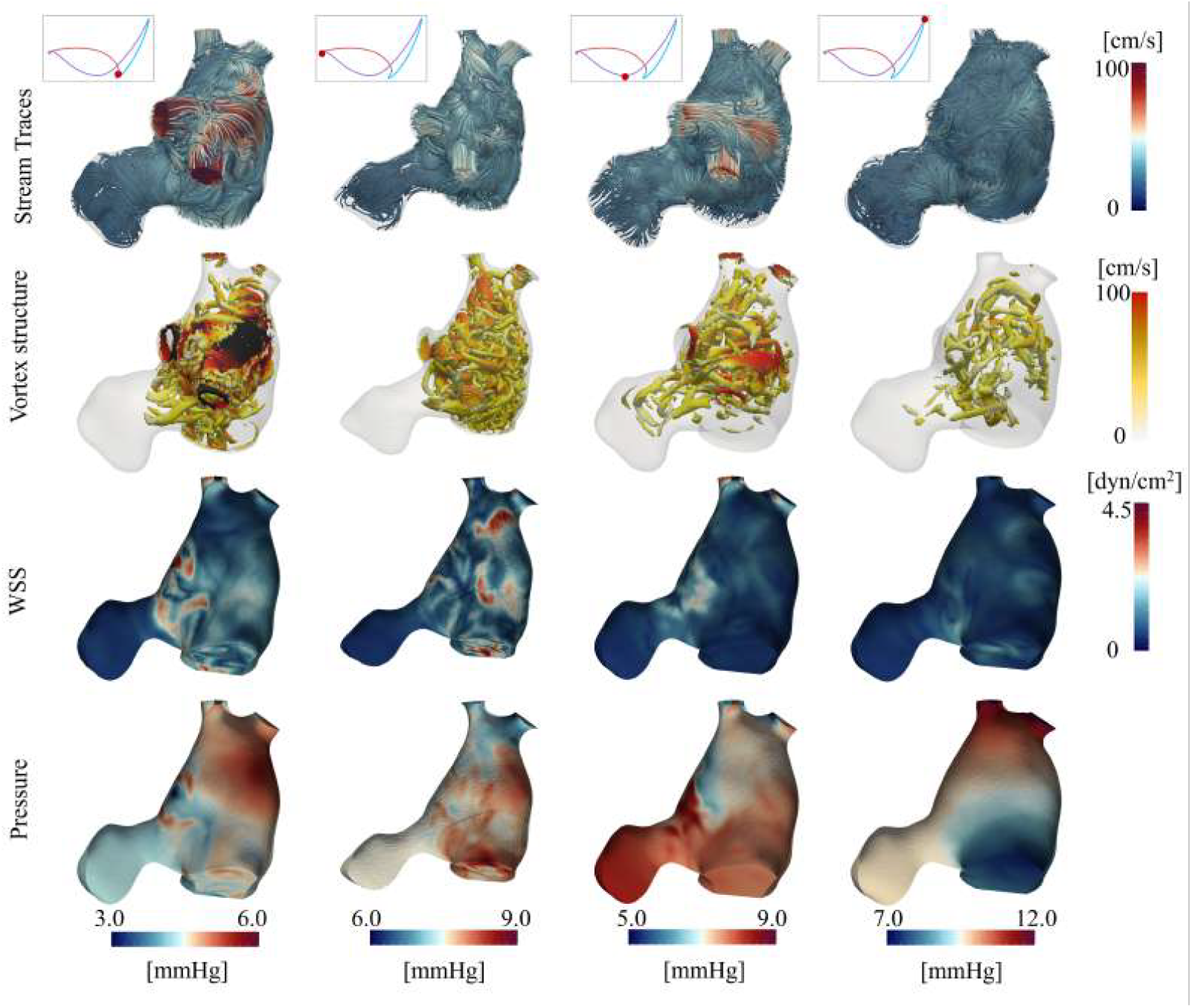
Characterizing LA hemodynamics along the anterior view during the cardiac cycle, simulated using personalized boundary conditions and wall motion extracted from the personalized, multiscale FEA modeling framework (Fig. 2). The complementary posterior view is included in Appendix C. (row-wise top to bottom) stream traces, vortex structures (identified using Q-criterion [136]) colored by velocity magnitude, wall shear stress (WSS), and surface pressure. (column-wise left to right) beginning of contraction or end of conduit phase, end of contraction (booster pump), mid-expansion (reservoir), and end of expansion or beginning of the conduit phase.

WSS, a key mechanobiological regulator of endothelial mechanosensing and a potential contributor to thrombogenesis in abnormally beating LA, is found to be highly dynamic and spatially varying. For instance, at the beginning of contraction, elevated WSS is observed on the anterior wall near the LA roof proximal to LAA (Fig. 11, third row, first column), whereas the inferior wall exhibits a concentrated patch of high WSS spanning the roof to the septum (third row and first column of Fig. C.16 in Appendix C). However, toward the end of the contraction, this patterning quickly shifts so that the inferior wall has more or less uniform shear, while the anterior wall exhibits pockets of high WSS (third row, second column in Fig. 11 and Fig. C.16 in Appendix C). During expansion, WSS is generally low and spatially uniform, with minor concentrations at the center of the inferior wall (third row and third column of Fig. C.16 in Appendix C)) and the near the LA roof proximal to LAA. Throughout the cycle, WSS is generally kept low in LAA relative to the bulk of the LA cavity, with the throat region between LAA and LA experiencing marginally high WSS.

Surface pressure distribution results reveal a consistent pressure gradient of approximately 3–5 mmHg from the pulmonary veins to the mitral valve at all phases of the cardiac cycle (bottom row in Fig. 11 and Fig. C.16). However, the absolute pressure values on the inner wall vary depending on the cardiac phase, with higher cavity pressures realized during LA contraction and expansion compared to the conduit phase. Moreover, the pressure also exhibits wide spatial heterogeneity, with different segments (pulmonary veins, mitral valve, and LAA) experiencing a cycle of high and low pressures.

## 4. Discussion

We developed a comprehensive workflow for simulating LA myocardial mechanics and blood flow using personalized tissue parameters (Fig. 2). The digital twinning framework integrates a patient’s time-resolved 3D CTA images and ECG data with a multiscale FEA model, created using our previously established segmentation pipeline [85], and a multi-step personalization workflow to closely align model-predicted deformation against image data. This last step is particularly a vital and novel aspect of the workflow, ensuring a reasonable replication of the patient’s physiology, while applying patient-specific boundary conditions and characterizing myocardial properties during both passive expansion and active contraction. By adopting a multiscale model setting, in which the 3D FEA model is coupled to a 0D LPN model, the current study overcomes one of the limitations in our earlier work on personalizing myocardial passive mechanics using iFEA [85]. In particular, instead of a generic pressure profile, the pressure applied during iFEA to estimate passive myocardial material parameters is derived from a surrogate-based LPN network with parameters fit to match clinical data.

Our model successfully replicates the patient’s physiology by matching against various clinical data, including cuff pressures, min-max cavity volumes of the four cardiac chambers, and other literature-based hemodynamic pressures (Fig. 7A). The 0D-based pressure profile extracted during passive expansion for passive mechanics characterization using iFEA results in a strong agreement between simulations and image-based volumes (Fig. 7C). Finally, tuning the active contraction parameters in a multi-scale setting captures the LA’s characteristic P–V diagram with both *a*– and *v* –loops (Fig. 8B). At the same time, global and local deformation also agree with data from time-resolved 3D CTA. This includes a reasonable agreement between LA and LAA volume changes throughout the cardiac cycle (Fig. 8C) and a closely aligned inner wall profile along an oblique plane at selected phases during the cycle (Fig. 8D).

The simulation-predicted myocardial motion, coupled with optimized pressures at pulmonary veins and mitral flow rate as boundary conditions, was then used to drive blood flow through the cavity. This tight integration between the clinical data and the multiscale models of myocardial mechanics and moving-domain blood flow will enable us to study the interplay between structural, geometric, and biomechanical factors elevating the risk of thrombosis and stroke under AF conditions. When coupled with the myocyte excitation models [137], the framework will allow us to assess stroke risk by predicting arrhythmogenic tissue deformations leading to altered blood flow and clot genesis, and analyze the interactions leading to clot fragmentation. Insights developed could potentially lead to personalized targeted treatment plans and mitigate the risk of device failure and anticoagulant overuse.

Below, we will discuss the rationale behind some of the modeling choices adopted in our approach, analyze the model’s sensitivities to key numerical parameters, and discuss limitations in the current work.

### 4.1. Rationale behind modeling choices

In developing the current modeling framework, we have made several modeling decisions related to model setup and methods employed, including geometry, material parameters, boundary conditions, and solver formulation. Here, we will discuss these choices, provide a rationale for each decision, and, where applicable, outline a procedure for improvement in the future.

The heterogeneity in LA wall thickness is well characterized in previous studies based on gross anatomical studies [39, 40] and MRI techniques [46]. This local variation in wall thickness and curvature is shown to impact regional wall stress distribution [49, 50]. LA, however, is relatively thin compared to the ventricular wall and is challenging to delineate its thickness on CTA data. Therefore, based on previously published data [52, 53, 55], we employed a uniform thickness of 2 mm in this work [56, 88, 91]. We will adapt our model creation workflow to account for variable wall thickness using methods tested for CTA images [48, 51].

Fiber-based constitutive models are widely adopted for LA mechanics [32, 50, 53, 56]. However, several studies have assumed transverse isotropy [50, 53, 56], partly informed by the experimental data [138]. In the current study, however, we adopted the orthotropic HO constitutive model to characterize the myocardial stress-strain relationship similar to the ventricles, as done in prior studies [28, 32, 85].

Likewise, although different methods were developed to represent LA fibers, including histology-based fiber rules [53] and DTMRI-guided rule-based methods [48, 88, 139], we employed a Laplace-Dirichlet-Rule-Based (LDRB) method developed by Piersanti et al. [88] to generate LA fiber and sheet orientations used in the constitutive model. Moreover, like most studies on LA mechanics cited above, we assumed the material parameters to be homogenous across the entire tissue, despite experimental evidence suggesting regional variation in LA tissue properties [138]. We will explore local material heterogeneity in a follow-up study.

To model the effects of the pericardium, we applied Robin boundary conditions on the epicardial surface (Eq. 7) to avoid the cost of contact mechanics based on the work of Pfaller et al. [32]. However, following the original work, we assumed these Robin parameters (stiffness and damping coefficients) to be spatially uniform [32]. Later simulation studies demonstrated that employing spatially varying epicardial boundary conditions may lead to better agreement with the image data [82, 83]. Although this spatial variation in Robin conditions seems vital for matching the ventricular deformation with image data, their role in aligning LA deformation is unclear. Nevertheless, we will explore this in a future investigation.

In our multiscale mechanics model, the applied hemodynamic pressure is time-dependent and coupled to the 0D LPN model in a closed-loop fashion. However, we assumed this pressure to be spatially uniform across the inner wall, which is the most common approach to model LA mechanics. However, our blood flow simulations suggest that the LA pressure is highly non-uniform. Together with the heterogeneity in geometry, material parameters, and epicardial boundary conditions, the non-uniform pressure can impact simulated deformation and wall stresses and strains. Further, although the LA pressures from the fully 0D model lie within the physiological range for a healthy subject [61], the P–V diagram is not physiological as it does not capture the *v*-loop (Fig. 7B). This is a limitation of our LPN model as it employs a linear pressure–volume relationship during the reservoir and conduit phases of the cycle and could be addressed using nonlinear models [61, 140, 141].

Although our motivation to develop the current framework for personalizing LA mechanics and blood flow is to study the stroke risk under AF conditions, the model cannot resolve myocyte ion exchange processes and is, therefore, not yet capable of predicting abnormal excitation and contraction patterns. Instead, we employed an active stress-based phenomenological model of contraction where the entire tissue contracts at once [32]. In a follow-up study, we will extend our modeling framework to account for LA myocyte excitation [137] and a realistic contraction model based on sarcomere kinematics [142]. We will tune the electrical conduction properties to match the patient’s ECG data (e.g., P-wave duration).

Conventional approaches to modeling incompressible solid mechanics typically treat pressure as a Lagrangian multiplier to enforce incompressibility as a constraint while the displacements are the primary unknown degrees of freedom [143]. In our approach, we treat pressure as the thermodynamic intensive state variable and it is equivalent to the mechanical pressure (i.e., hydrostatic or dilatational part of the Cauchy stress, Eqs. 6b,c), invoking the Stokes hypothesis (see Sections 2.2–2.4 in Liu & Marsden [93]). As a result, the usually adopted Helmholtz free energy for the volumetric penalization, written as a function of the Jacobian (*J*, Eq. 2), is transformed to Gibbs free energy, a function of the thermodynamic pressure, via a Legendre transformation as pressure and specific volume are work conjugates [144]. The motivation to use pressure instead of *J* for volume-preserving free energy (Gibbs vs. Helmholtz) in the incompressible limit (i.e., *J* = 1) is the degeneracy of pressure computed as a derivative with respect to *J*, which is a constant, and this definition becomes invalid (see Remarks 4 and 6 in Liu & Marsden [93]). On the contrary, employing velocity and pressure as the unknown degrees of freedom will transform the algebraic constraint for mass conservation into a PDE (i.e., divergence-free velocity in the incompressible limit, Eq. 5b), similar to the Navier-Stokes equations for fluid flow (Eq. 14), and leads to a well-defined formulation for both compressible and incompressible materials. Although both these approaches are implemented in our solver and have been verified for cardiac mechanics [95], we chose to employ a velocity-pressure-based formulation (Eqs. 5–7) in this work for the arguments presented above.

### 4.2. Sensitivity to mesh size

We evaluated the sensitivity of the LA mechanics model to mesh edge size. In particular, we varied the mesh element size and examined its impact on the LA P–V diagram (Fig. 12). We tested four different element sizes that systematically refined the number of layers in the transmural direction, from a single layer to four layers across the myocardial thickness (Fig. 12A).

**Figure 12:**
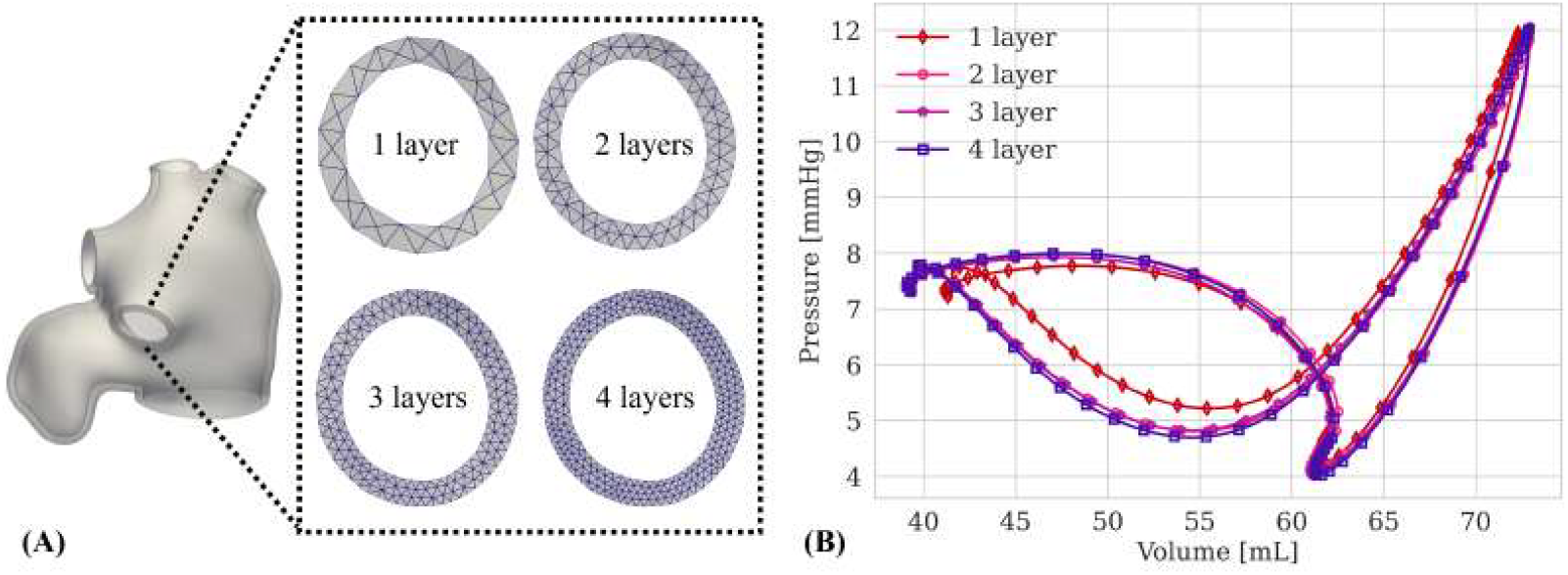
Mesh convergence analysis. (A) LA mesh is systematically refined along the transmural direction by increasing the number of layers across the thickness. (B) Comparison of LA pressure–volume (P–V) loops for different mesh sizes.

This resulted in approximately 11K, 90K, 320K, and 880K elements across the entire tissue. All other simulation parameters, including LPN, material, and boundary conditions, were kept the same (Section 3.2).

Each simulation was run for 5 cycles until it reached a limit cycle with negligible cycle-to-cycle variations. The total CPU time increased substantially with successive mesh refinement. While the single-layer mesh case was run on a local workstation with an Intel-14900 5.8GHz CPU with 24 cores, all other cases were simulated on Columbia’s Ginsburg computing cluster, with each node containing an Intel Xeon Gold 6226 2.9GHz CPU with 32 cores. The simulation run times were 120s, 1300s, 3700s, and 10,500s for the single-layered, two-layered, three-layered, and four-layered meshes, respectively. While the two-layered case was run using a single node on the cluster, the three- and four-layered cases were run using three nodes on the same computing cluster.

Except for the single-layered mesh, all other meshes showed only subtle differences in the P–V diagram (Fig. 12B). Therefore, to achieve a balance between accuracy and computational efficiency, we selected the two-layered mesh for iFEA, as it involves hundreds of iterations during the optimization process, and the four-layered mesh for the multiscale mechanics simulation, as the resulting inner wall deformation from this multiscale simulation is used as input to drive blood flow.

### 4.3. Sensitivity to active stress magnitude

In this study, we employed a phenomenological model of the active stress (Eq. 9) to simulate LA contraction, in which the contractility parameter, *σ*_0_, controls the magnitude of the peak active stress. We analyzed the sensitivity of the LA P–V diagram by perturbing *σ*_0_ by 5% around the baseline value of 36 kPa (Eq. 9A). It is not surprising that small perturbations in the contractility proportionally changed the minimum volume attained during the LA booster phase without affecting the pressures, with lower contractility leading to a slightly higher minimum volume and vice versa. It is also interesting to note that these small perturbations in *σ*_0_ did not have any noticeable impact on the conduit phase.

### 4.4. Sensitivity to fiber directions

In this study, we created LA myofibers using a procedure outlined in Piersanti et al. [88]. The fiber directions are parameterized using a region-specific parameter, *τ*, and the baseline values are provided in Section 2.2.1 for LPV, RPV, and MV. Here, we simultaneously varied all these values by ± 30% of the baseline value to evaluate the mechanics model’s sensitivity to the fiber directions.

Our results suggest that the LA P–V diagram is less impacted by the changes in fiber directions (Fig. 14). The overall magnitudes of the fiber stretch remain the same for all the cases, and the histogram distributions are only marginally affected by the *τ* parameter (Fig. C.18 in Sec. Appendix C).

**Figure 13:**
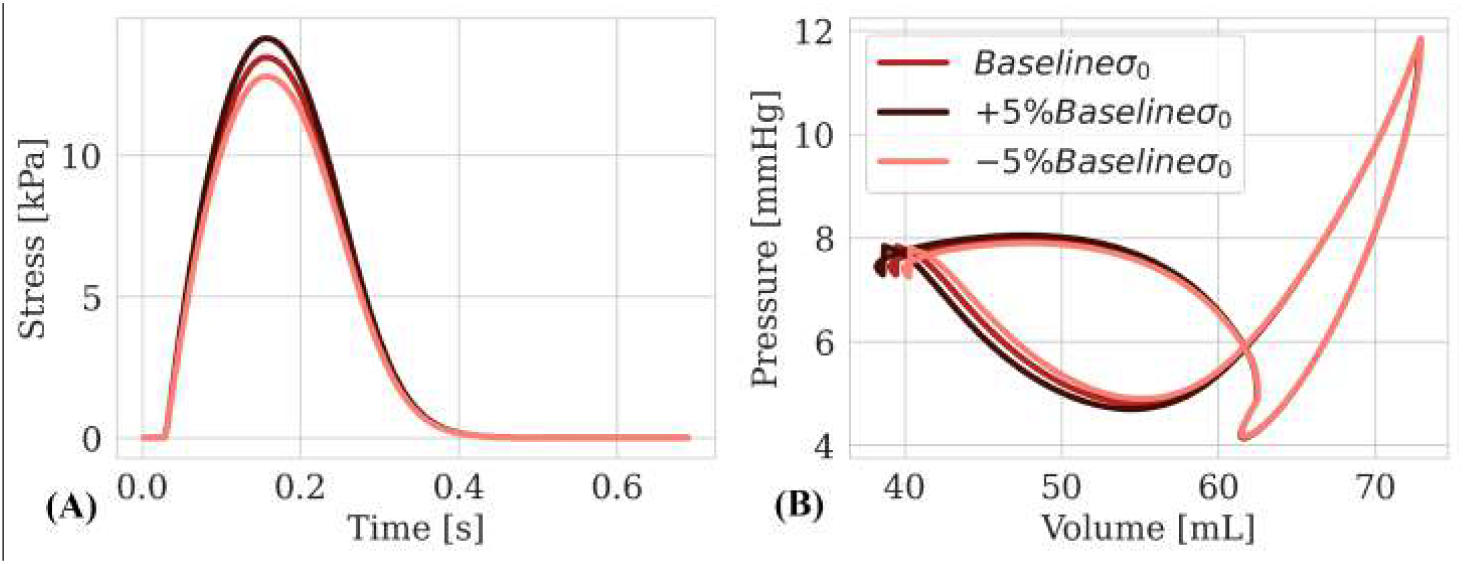
Sensitivity to contractility parameter, *σ*_0_ (Eq. 9). (A) Active stress profiles for the baseline contractility (*σ*_0_ = 36 *kPa*) and ± 5% variation. (B) Comparison of the corresponding pressure–volume (P–V) diagrams for different contractility parameters.

**Figure 14:**
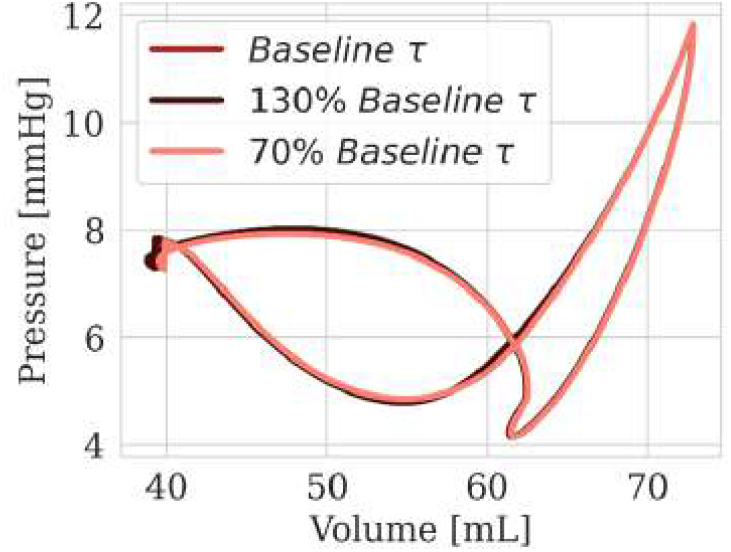
Comparison of the left atrial (LA) pressure–volume (P–V) diagram for ± 30% variations in the baseline region-specific fiber direction parameters (*τ*_lpv_, *τ*_rpv_, and *τ*_mv_).

However, the spatial distribution of the local fiber stretch suggests loose dependence on the fiber parameter, especially on the mid-anterior wall when the fibers sharply change directions and the location of this sudden inflection is dependent on *τ* (Fig. C.17 in Appendix C).

### 4.5. Limitations and future perspectives

Our work has several limitations. Many of these limitations stem from the choices we adopted in setting up the FEA model, including geometric and material considerations, boundary conditions, and the physical processes resolved. These have been discussed in Section 4.1, along with our plan to address them in the future. Additionally, we tested our model only on one patient. In the future, we plan to apply our model to a larger and diverse patient cohort, spanning age groups, sex, and varied LA sizes and systemic hemodynamic conditions. We will also plan to assess our model’s sensitivity to assumptions on the EMD duration used for computing activation start and end times from the ECG data (Appendix B). The effects of literature-derived pressures (e.g., min/max pulmonary artery pressures, PAWP, and CVP in Table 2) for personalizing the 0D LPN model on the estimated tissue parameters and overall mechanical deformation will be evaluated as well. To limit the overall computational time for characterizing the passive mechanics, we will explore employing data-driven models, particularly neural network-based emulators to directly predict myocardial deformation given a geometry, material parameters, and boundary conditions [145, 146].

## 5. Conclusion

In this study, we introduced a novel digital twinning framework that integrates 3D myocardial mechanics with a 0D lumped parameter network (LPN)-based blood flow model to simulate patient-specific wall deformation and hemodynamics in the left atrium (LA). By leveraging time-resolved 3D CTA images and clinical measurements, including ECG readings and cuff pressures, we reconstructed the LA myocardium, optimized the 0D model parameter to fit clinical measurements, and characterized the tissue’s passive material properties through an inverse finite element analysis (iFEA). The multiscale coupling between the 3D mechanics model and the 0D LPN model effectively captures the interaction between LA mechanics and the cardiovascular system, reproducing the classic 8-shaped pressure–volume diagram. The active contraction model parameters are further tuned to closely align with CTA-based volumes and local deformation. The resulting simulation-predicted, personalized endocardial wall motion, together with pulmonary venous pressures and mitral flow rate from the 0D LPN model, are prescribed to drive blood flow through the LA cavity. Ultimately, this study lays the groundwork for creating an end-to-end multiphysics modeling framework to advance our understanding of the LA structure-function relationships contributing to elevated stroke risk from atrial fibrillation, which affects millions of people worldwide.

## Acknowledgments

The authors would like to acknowledge financial support from the American Heart Association’s Second Century Early Faculty Independence Award (#24SCEFIA1260268) in performing this work. The authors would also like to acknowledge computing resources and services received through the

Columbia Shared Research Computing Facility and the Ginsburg High-Performance Computing Cluster.

## Appendix A. LPN model equations

We employed a simplified LPN model of the circulatory system adapted from prior work [86, 87]. As discussed in Section 2.2.3, our compartmental LPN model comprises individual blocks for RA, RV, and LV, modeled using time-varying elastance function, while the systemic and pulmonary arterial and venous blocks are modeled using R-L-C circuit elements (Fig. 4). Below, we describe the governing differential algebraic equations (DAEs) of the full LPN model, incorporating a 0D surrogate model for LA. In the following subsection, we will describe the changes to these equations for coupling with the 3D FE model.

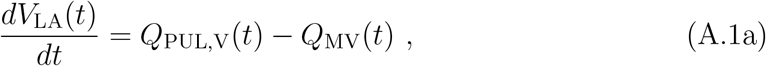

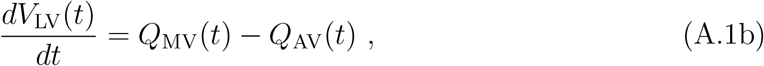

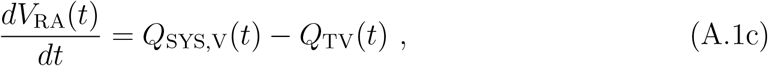

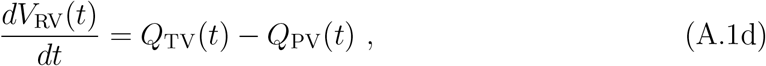

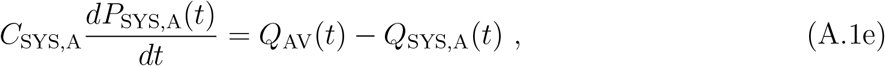

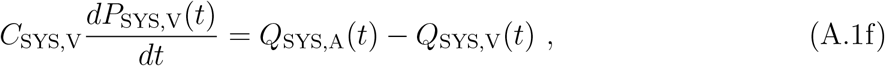

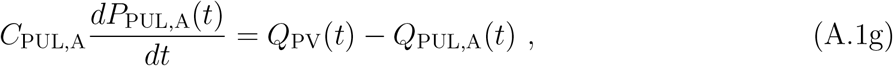

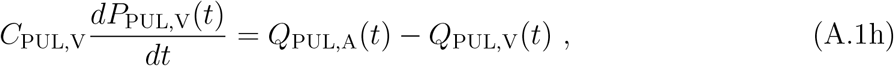

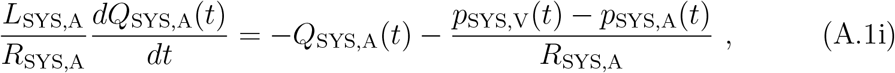

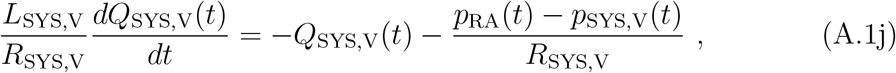

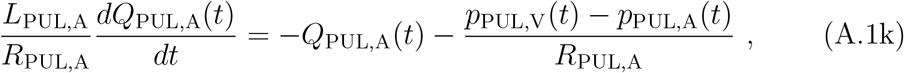

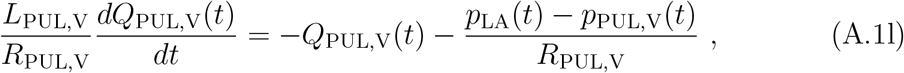

where, *Q, P*, and *V* represent the flowrate, pressure, and volume, respectively, while *R, C*, and *L* denote the resistance, capacitance, and inductance, respectively. The subscripts LA, LV, RA, and RV represent the left atrium, left ventricle, right atrium, and right ventricle, respectively, whereas the subscripts PUL and SYS denote the corresponding values for the pulmonary and systemic circulations, and A and V represent the corresponding values for arterial and venous circulations. The valvular flowrates are related to pressure gradients across the valves as,

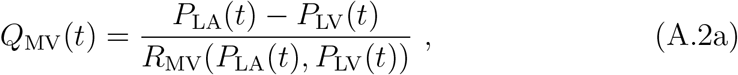

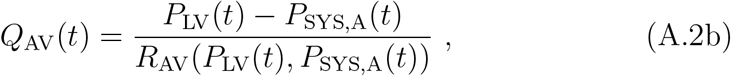

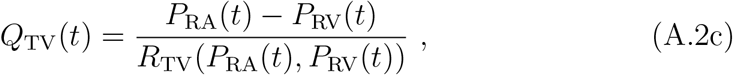

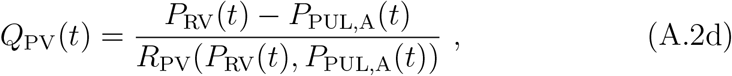

where,

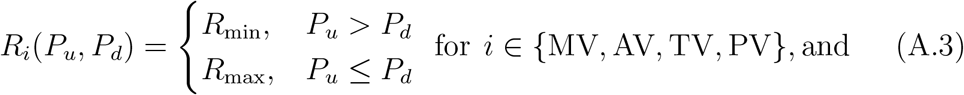

*P*_*u*_ denotes the pressure upstream of the valve, and *P*_*d*_ is the downstream pressure. The chamber pressures are calculated as,

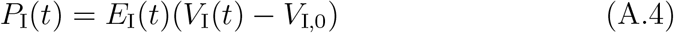

where *V*_I_(*t*) is the volume of chamber I **∈** {LA, RA, LV, RV}, *V*_I,0_ is the resting volume of chamber I, and the time-varying elastance *E*_I_(*t*) is given by,

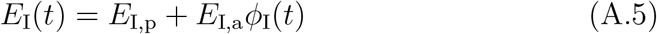

where *E*_I,p_ governs the passive stiffness and *E*_I,a_ drives the contraction with a time-dependent activation function, *ϕ*(*t*). In this work, we adopted a double-Hill activation profile to model the chamber contraction given by [114],

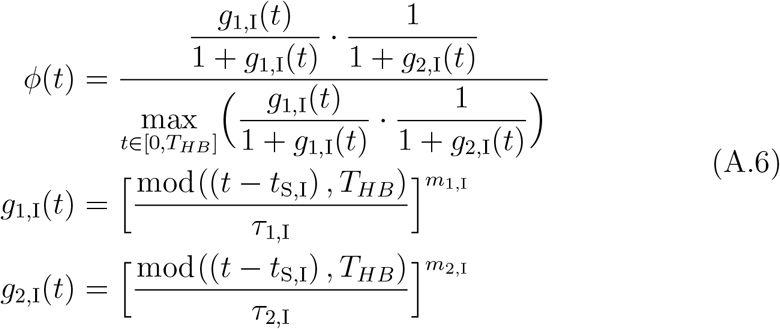

where *t*_S,I_ is the start time of contraction, *τ*_1,I_, *τ*_2,I_ control the time shift, and *m*_1,I_, *m*_2,I_ control the slope of each Hill function [114], defined for each chamber I **∈** {LA, LV, RA, RV}, and *T*_*HB*_ is the duration of one cardiac cycle (*T*_*HB*_ = 60*/HR*). Our approach to setting these parameters is provided in Appendix B.

### Appendix A.1. Revised LPN equations for coupled 0D/3D model

The DAEs described in the previous Section Appendix A correspond to the full 0D LPN model, assuming a surrogate model for the LA. Here, the volume change of LA is governed by the net flow across it (Eq. (A.1b)), and the cavity pressure is related to its volume using a time-varying elastance model (Eq. (A.4)). However, when the surrogate model is replaced by the 3D FE model, the flowrate (*Q*_LA_) is passed from the 3D solver to the 0D solver, and the 0D-computed pressure (*P*_LA_) is transferred back to the 3D solver (Fig. 4). Consequently, the LA volume rate equation (Eq. (A.1b)) is replaced by an algebraic constraint, which in turn informs the LA hemodynamic pressure calculation as,

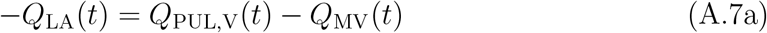

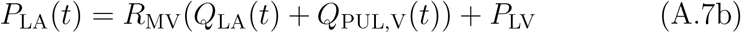

As noted in Section 2.2.3, for every Newton iteration of the 3D solver, the 0D LPN model is integrated from timestep *n* to *n* + 1. This makes the above Eq. (A.7) an implicit equation for 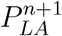 as the mitral flow and valve resistance depend on the pressure difference between LA and LV (Eq. (A.2b)). To avoid numerical issues associated with valve opening and closing between Newton iterations, we make Eq. (A.7) explicit and use 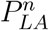 to evaluate the state of the mitral valve and determine its resistance and flowrate through it [86].

## Appendix B. ECG-based activation times

Key measures extracted from the patient’s ECG include the PR, QRS, and QT duration intervals (Table 1). The PR interval is associated with atrial depolarization and the conduction time from the atria to the ventricles (Fig. 2, left). The electrical activation of the ventricle or the ventricular depolarization is characterized by the QRS complex [147], while the T-wave is associated with ventricular repolarization, reflecting the recovery phase of the ventricles [148, 149, 150]. Assuming that the chamber’s mechanical contraction has a delayed start, quantified by the electromechanical delay (EMD), which is estimated to be 25 ms [151], we can determine the contraction start and end times of the atria and ventricles as,

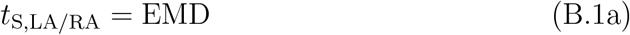

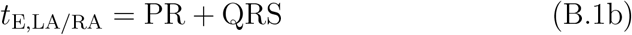

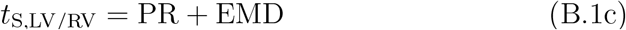

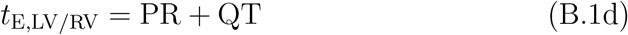

where *t*_S_ and *t*_E_ represents the start and end times of chamber contraction. Further, we assumed the P-wave onset occurs at *t* = 0 ms and ignored any activation onset delay between the two atria and ventricles.

**Table B.6:** Contraction start and end times and other parameters of the double Hill function.

| Chamber | $t_S$ (s) | $t_E$ (s) | $\tau_1$ (s) | $\tau_2$ (s) | $m_1$ | $m_2$ |
| --- | --- | --- | --- | --- | --- | --- |
| LA | 0.025 | 0.270 | 0.086 | 0.148 | 1.32 | 13.1 |
| RA | 0.025 | 0.270 | 0.086 | 0.148 | 1.32 | 13.1 |
| LV | 0.207 | 0.566 | 0.297 | 0.387 | 1.32 | 27.4 |
| RV | 0.207 | 0.566 | 0.297 | 0.387 | 1.32 | 27.4 |

The time shift parameters (*τ*_1_, *τ*_2_) of the double Hill activation function are computed as,

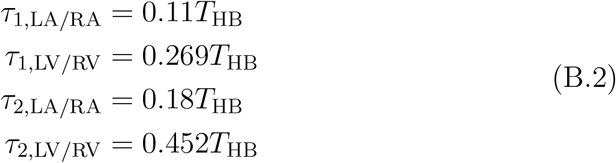

while slope parameters (*m*_1_, *m*_2_) are extracted from prior work [114]. Table B.6 summarizes the contraction start and the double Hill activation function parameters for all cardiac chambers.

## Appendix C. Additional data from multiscale mechanics and blood flow simulations

Figs. C.15 and C.16 provide complementary views of the LA myocardial mechanics and blood flow simulation data, described in Sections 3.3 and 3.4 of the main manuscript. Fig. C.17 compares fiber stretch during the cardiac cycle for ± 30% variations in the baseline region-specific fiber direction parameters (*τ*_lpv_, *τ*_rpv_, and *τ*_mv_ provided in Section 2.2.1).

**Figure C.15:**
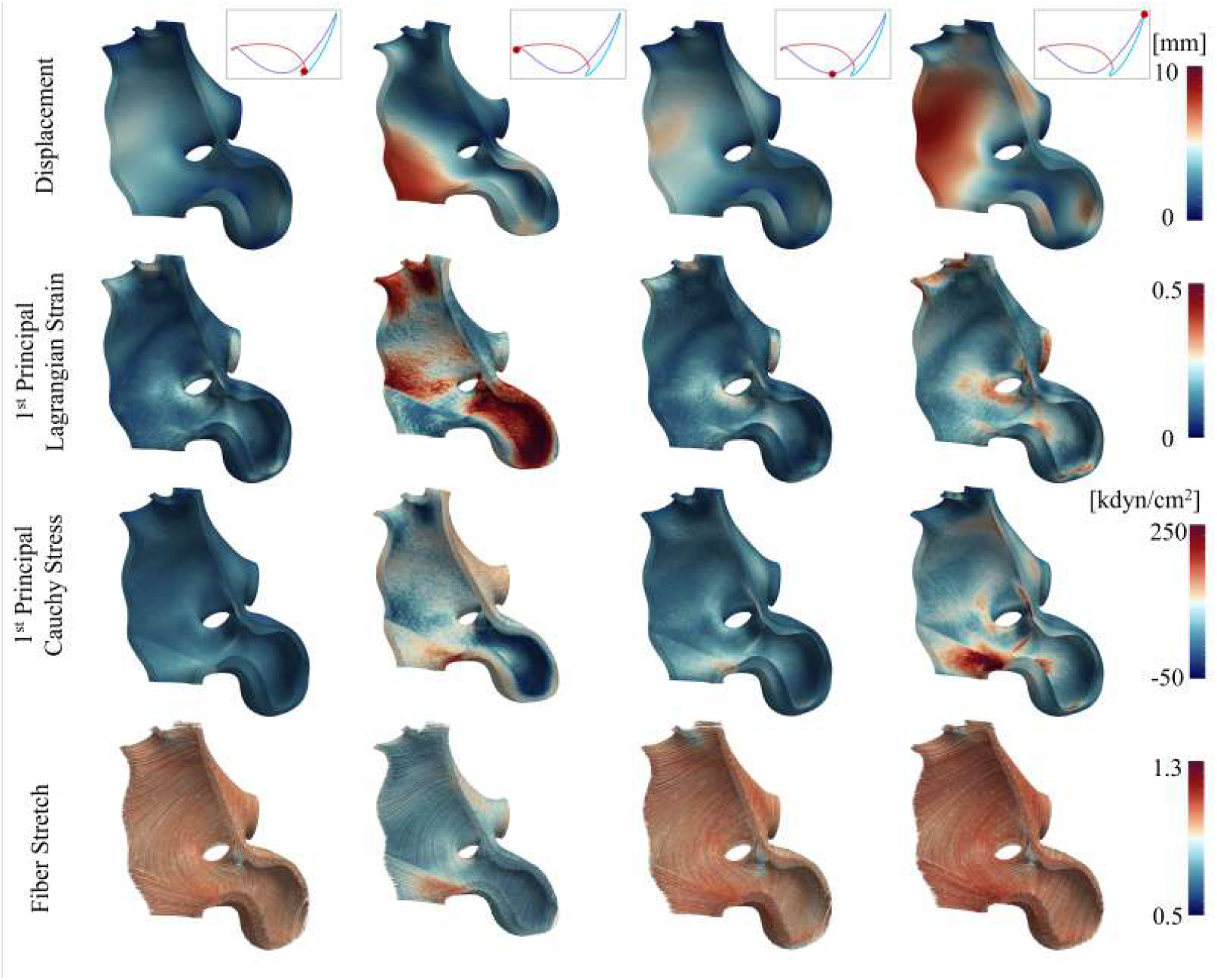
Characterizing LA mechanics on the inferior wall viewed along the posterior-anterior perspective during the cardiac cycle using the current personalized, multiscale FEA modeling framework (Fig. 2). The complementary anterior clip is shown in Fig. 10 of the main manuscript. (row-wise top to bottom) displacement, 1^st^ principal (Green-) Lagrange strain, 1^st^ principal Cauchy stress, and fiber stretch. (column-wise left to right) beginning of contraction or end of conduit phase, end of contraction (booster pump), mid-expansion (reservoir), and end of expansion or beginning of the conduit phase.

**Figure C.16:**
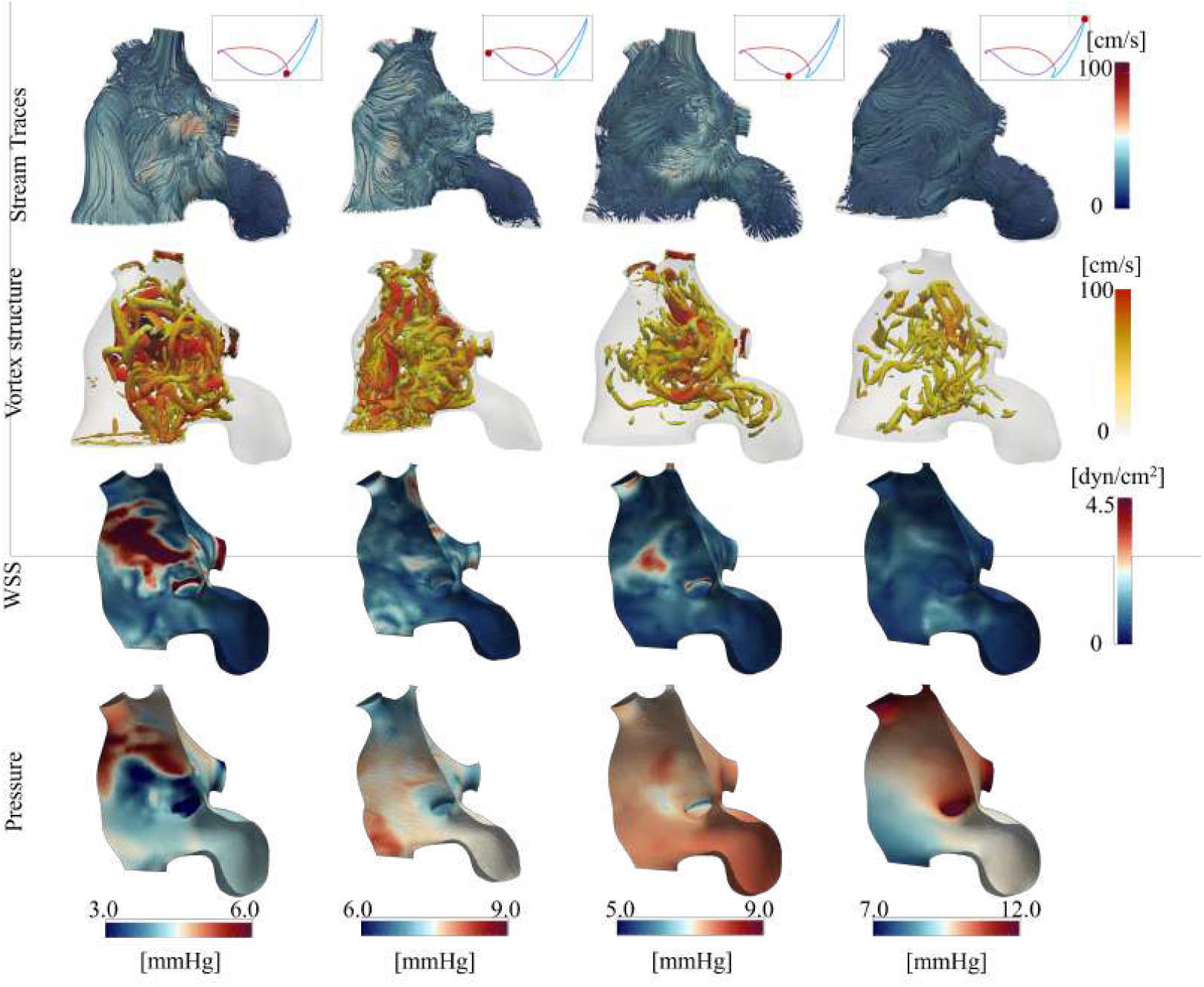
Characterizing LA hemodynamics along the posterior view during the cardiac cycle, simulated using personalized boundary conditions and wall motion extracted from the current personalized, multiscale FEA modeling framework (Fig. 2). The anterior view of the same is included in Fig. 11 of the main manuscript. (row-wise top to bottom) stream traces, vortex structures (identified using Q-criterion [136]) colored by velocity magnitude, wall shear stress (WSS), and surface pressure. (column-wise left to right) beginning of contraction or end of conduit phase, end of contraction (booster pump), mid-expansion (reservoir), and end of expansion or beginning of the conduit phase.

**Figure C.17:**
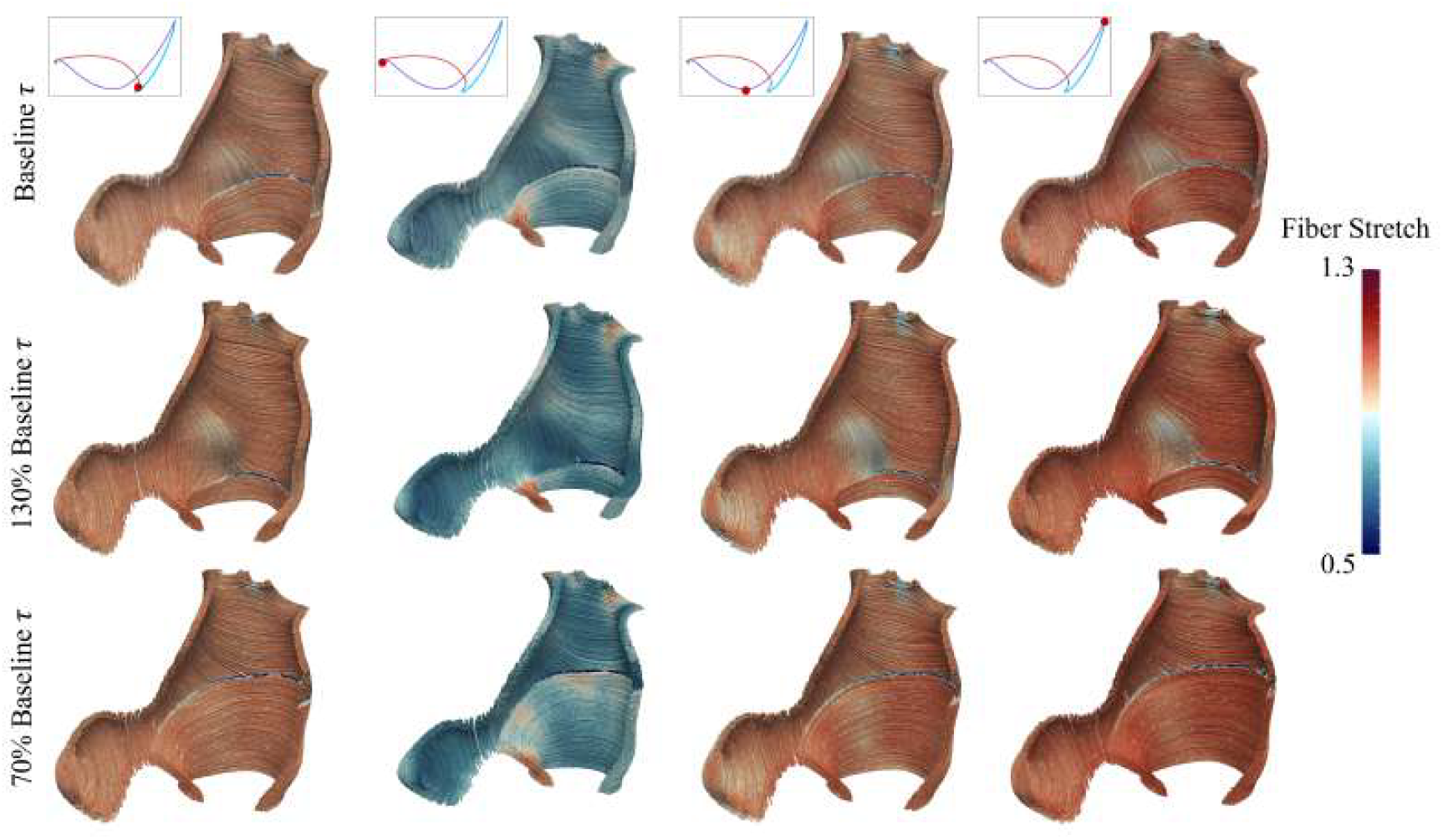
Comparison of the left atrial (LA) fiber stretches along the anterior view at select phases during the cardiac cycle for ± 30% variations in the baseline region-specific fiber direction parameters (*τ*_lpv_, *τ*_rpv_, and *τ*_mv_). (row-wise top to bottom) baseline *τ*, +30% baseline *τ*, and − 30% baseline *τ*. (column-wise left to right) beginning of contraction or end of conduit phase, end of contraction (booster pump), mid-expansion (reservoir), and end of expansion or beginning of the conduit phase.

**Figure C.18:**
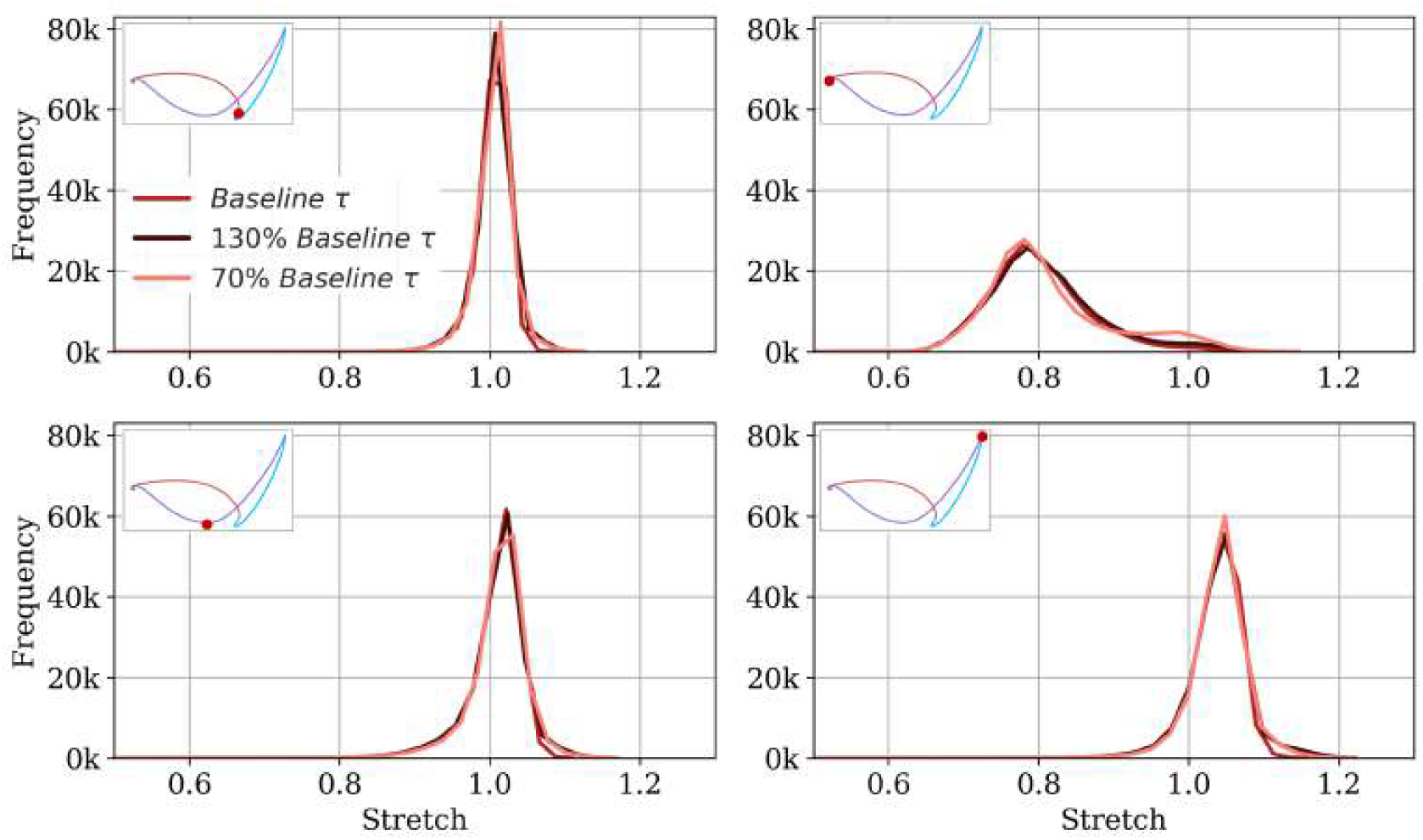
Comparison of the histogram of left atrial (LA) fiber stretch distribution for ± 30% variations in the baseline region-specific fiber direction parameters (*τ*_lpv_, *τ*_rpv_, and *τ*_mv_).

## Footnotes

* Our choice of 20% R-R as the starting point is founded on the rationale that LA tissue is fully contracted at the beginning of systole (0% R-R) from the preceding atrial kick (booster pump). We, therefore, allow the LA to relax due to the elastic recoil effect during early systole that manifests as a drop in the cavity pressure with a concomitant increase in volume. We characterize passive mechanics when the cavity pressure and volume are directly proportional to each other during the reservoir phase (Fig. 1B) [85].

† https://simvascular.github.io/

‡ Autodesk Inc., https://www.meshmixer.com/, version: 3.5

§ https://github.com/neka-nat/probreg

¶ https://wias-berlin.de/software/tetgen/

‖ https://simvascular.github.io/

** https://github.com/SimVascular/svFSI

†† https://trilinos.github.io/

* https://wias-berlin.de/software/tetgen/

† https://simvascular.github.io/

